# Paclitaxel compromises nuclear integrity in interphase through SUN2-mediated cytoskeletal coupling

**DOI:** 10.1101/2025.01.17.633376

**Authors:** Thomas Hale, Victoria L Hale, Piotr Kolata, Ália dos Santos, Matteo Allegretti

## Abstract

Regulation of Lamin A/C levels and distribution is crucial for nuclear integrity and mechanotransduction via the Linker of Nucleoskeleton and Cytoskeleton (LINC) complex. Dysregulation of Lamin A/C correlates with poor cancer prognosis, and its levels determine sensitivity to the microtubule-stabilising drug paclitaxel. Paclitaxel is well-known for disrupting mitosis, yet it also reduces tumour size in slow-dividing tumours, indicating an additional, poorly characterised interphase mechanism.

Here, we reveal that paclitaxel induces nuclear aberrations in interphase through SUN2-dependent Lamin A/C disruption. Using advanced optical imaging and electron cryo-tomography, we show the formation of aberrant microtubule-vimentin bundles during paclitaxel treatment, which coincides with nuclear deformation and altered Lamin A/C protein levels and organisation at the nuclear envelope. SUN2 is required for Lamin A/C reduction in paclitaxel and is in turn regulated by polyubiquitination. Furthermore, Lamin A/C expression levels determine not only cell survival during treatment but also recovery after drug removal.

Our findings support a model in which paclitaxel acts through both defective mitosis and interphase nuclear-cytoskeletal disruption, providing additional mechanistic insights into a widely used anticancer drug.

## Introduction

Paclitaxel is a taxane widely used to treat breast, ovarian and lung cancers^1–3^. It acts by binding stoichiometrically to β-tubulin within microtubules and inducing conformational changes that stabilise interactions between adjacent α/β-tubulin heterodimers^4^, preventing microtubule depolymerisation and enhancing bundle formation^5,6^. Historically, paclitaxel’s anti-cancer mechanism was thought to rely on its suppression of microtubule dynamics in the mitotic spindle, which activates the mitotic checkpoint to result in cell cycle arrest and subsequently apoptosis^7^. Since then, it has been shown that cells can escape mitotic arrest in clinically relevant concentrations of paclitaxel, resulting in multipolar divisions due to improper chromosome segregation, ultimately leading to cell death.^8,9^ However, paclitaxel reduces tumour size even in tumours with low duplication rates and where only a small fraction of cells is proliferative^10–12^. Furthermore, intravital imaging of tumours in mice treated with taxanes shows that cell death is predominantly induced independently of mitotic defects^11^. This suggests that paclitaxel’s anti-cancer mechanism likely involves an additional activity in interphase^13–15^.

There have been several hypotheses proposing how paclitaxel may induce apoptosis in interphase, including the activation of pro-apoptotic signalling following the disruption of microtubule structure^16,17^, and perturbation to mitochondria^18,19^. Since microtubules perform myriad functions in the cell, it is likely that perturbations to multiple different aspects of cellular function contribute to apoptosis. Recently, Smith *et al* (2021) showed that paclitaxel treatment leads to fragmentation of the nucleus independently of cell division^14,20,21^. Together with observations that paclitaxel results in microtubule re-organisation around the nucleus during interphase^22,23^ and ectopic localisation of nuclear envelope (NE) and lamina proteins in cancer cells^24^, this suggests that an additional anti-cancer mechanism of paclitaxel may be via the disruption of nuclear-cytoskeletal coupling, leading to an increased risk of DNA damage and consequent cell death^25–27^. However, this mechanism is poorly understood.

Nuclear-cytoskeletal coupling is mediated by Linker of Nucleoskeleton and Cytoskeleton (LINC) complexes, which consist of a trimer of SUN proteins that span the inner nuclear membrane and interact at the C-terminal end of a periplasmic coiled-coil region with the C-terminus of three KASH protein monomers^28^. KASH proteins traverse the outer nuclear membrane and connect to the cytoskeleton in the cytoplasm at their N-terminus^29^, while the N-terminus of SUN proteins connects to the nuclear lamina in the nucleoplasm^30^. Cytoskeletal forces are therefore transmitted to the nuclear lamina and linked chromatin domains via these direct connections, which provide the physical basis for mechanotransduction^31^.

The nuclear lamina plays an important role in controlling the mechanical response of the nucleus, ensuring it can withstand and react to mechanical forces^32,33^. The nuclear lamina consists of the A-type lamins Lamin A and C, which are splice isoforms of the *LMNA* gene, as well as the B-type lamins Lamin B1 and B2^34^. These are intermediate filament proteins that polymerise to form a filamentous mesh beneath the NE^34^. Aberrations to the nuclear lamina and nuclear morphology are prevalent across many cancer types and are often used as biomarkers in diagnosis^35^. In particular, Lamin A/C expression is frequently altered in cancer cells^36–40^, which is thought to play a role in migration and invasion, a hallmark of cancer that leads to metastasis^40^. However, alterations to the nuclear lamina may also make cancer cells particularly sensitive to perturbations to nuclear-cytoskeletal coupling. In support of this, decreased Lamin A/C expression has been shown to increase sensitivity to paclitaxel in ovarian cancer cells^14^.

Here we propose that cancer cells have increased vulnerability to paclitaxel both during interphase and following aberrant mitosis due to pre-existing defects in their NE and nuclear lamina. Supporting this, we show that paclitaxel disrupted the organisation of microtubule, actin, and vimentin filaments around the nucleus during interphase leading to nuclear deformation. Nuclear deformation was more severe in cells with decreased levels of Lamin A/C. We also report similar nuclear deformation phenotypes in cells where microtubules were stabilised by overexpression of Tau, thus linking the observed nuclear deformation in paclitaxel with its microtubule bundling activity. Our data show that paclitaxel treatment severely affected nuclear lamina organisation and that this occurred via connection to SUN2-containing LINC complexes, which were regulated by ubiquitination following paclitaxel treatment. Finally, we show that Lamin A/C expression levels affected overall sensitivity to paclitaxel and, most importantly, recovery from it.

Altogether, our data provides insights into interphase mechanisms of paclitaxel and the role of the nuclear lamina in drug sensitivity.

## Results

### Paclitaxel induces cytoskeletal reorganisation around the nucleus during interphase

To investigate how paclitaxel affects the organisation of microtubules during interphase, human fibroblasts were treated with 5 nM paclitaxel or control media for 16 h. Experiments were performed at 5 nM paclitaxel (with additional experiments to determine dose relationships at 1 and 10 nM) because this aligns with previous studies^7,14,24^. Furthermore, previous analysis of patient plasma reveals that typical concentrations are within the low nanomolar range^8^, and concentrations of 5-10 nM are required in cell culture to reach the same intracellular concentrations observed *in vivo* in patient tumours^9^. This aligns with *in vitro* cytotoxic studies of paclitaxel in eight human tumour cell lines which show that paclitaxel’s IC50 ranges between 2.5 and 7.5 nM^41^.

Following paclitaxel treatment, cells were analysed by immunofluorescence confocal microscopy for α-tubulin and stained for DNA with Hoechst (Figure 1A). In contrast to control cells where microtubules were dispersed throughout the cytoplasm and emanated from a single microtubule organising centre (MTOC), in paclitaxel-treated cells the microtubules formed dense rings around the nucleus with no single obvious MTOC (Figure 1A). To determine if these microtubules surrounding the nucleus were bundled, we used 2D stochastic optical reconstruction microscopy (STORM) for high-resolution morphological characterisation (Figure 1B). Quantitative HDBSCAN clustering of STORM data revealed an increase of 42% in the microtubule bundle diameter in the presence of paclitaxel compared to untreated cells (Figure 1C).

**Figure 1.**
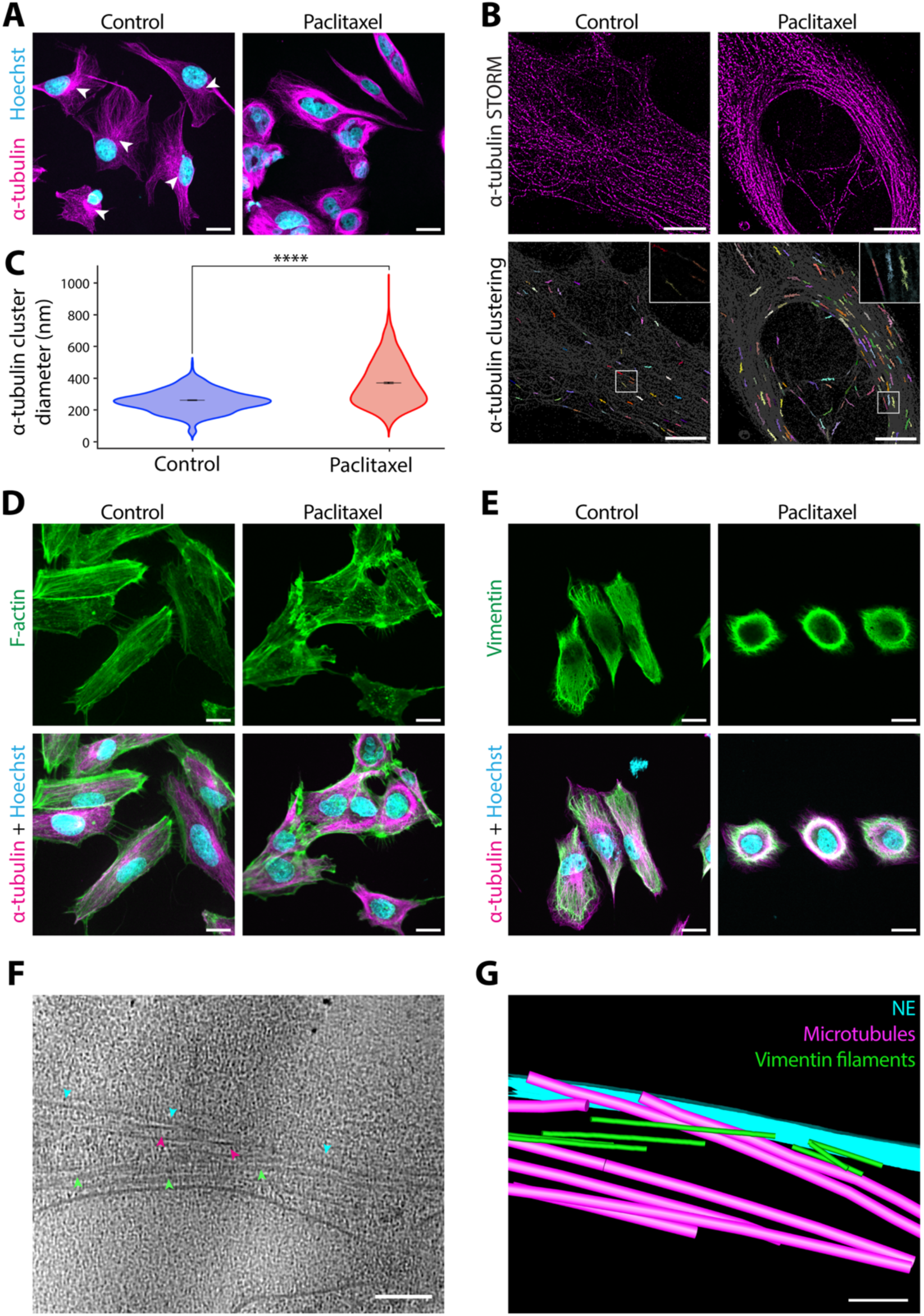
Paclitaxel-induced cytoskeletal reorganisation around the nucleus in interphase. **(A)** Confocal micrographs of cells fixed after 16 h incubation in control media or 5 nM paclitaxel. DNA was stained using Hoechst (cyan), and microtubules using α-tubulin immunofluorescence (magenta). MTOCs are marked with arrowheads. Scale bars = 20 μm. **(B)** STORM imaging of α-tubulin immunofluorescence in cells fixed after 16 h incubation in control media or 5 nM paclitaxel. Lower panels show α-tubulin clusters generated with HDBSCAN analysis. Different colours distinguish individual α-tubulin clusters, representing individual microtubule filaments or filament bundles. Scale bars = 10 μm. **(C)** Violin plot comparing the diameter of α-tubulin clusters from (B) between control cells (blue) and paclitaxel-treated cells (red). The mean ± the standard error of the mean (SEM) is shown in black: control = 260 ± 12.5 nm, n = 1116 across 20 cells; paclitaxel = 369 ± 17.9 nm, n = 1354 across 20 cells. Statistical analysis: t-test, P = 3.39 × 10^−16^ (****). **(D-E)** Confocal micrographs of control and paclitaxel-treated cells stained for DNA using Hoechst (cyan), microtubules using α-tubulin immunofluorescence (magenta), and either **(D)** F-actin using Alexa488-phalloidin (green), or **(E)** vimentin using immunofluorescence (green). Scale bars = 20 μm. **(F)** 2D slice of a reconstructed tomogram of the NE region in a paclitaxel-treated cell. Bundled vimentin filaments (green arrowheads) and microtubules (magenta arrowheads) are seen closely apposed to the NE (cyan arrowheads). Scale bar = 100 nm. **(G)** Segmentation of the NE (cyan), microtubules (magenta), and vimentin filaments (green) from the tomogram in (F). Scale bar = 100 nm.

Since there is extensive crosstalk between microtubule, actin, and intermediate filaments^42,43^, we next tested how the organisation of actin and vimentin was affected by paclitaxel treatment (Figure 1D and E). As expected, actin filaments were distributed throughout the cortex in control cells, but in the presence of paclitaxel, perinuclear actin was instead condensed into puncta (Figure 1D). Interestingly, vimentin colocalised strongly with microtubules^44,45^, and also reorganised into a dense network surrounding the nucleus following paclitaxel treatment (Figure 1E). The extent of the cytoskeletal reorganisation correlated with the concentration of paclitaxel, as microtubules became increasingly bundled and the actin cytoskeleton more condensed with increasing concentrations (0 to 10nM) of paclitaxel (Supplementary Figure 1A). Similar cytoskeletal reorganisation around the nucleus was also observed in human breast cancer MDA-MB-231 cells in 5 nM paclitaxel (Supplementary Figure 1B).

To visualise the rearranged cytoskeleton at high resolution and investigate how it interacts with the nucleus, we used electron cryo-tomography (cryo-ET). Cells were first thinned by cryo-focused ion beam (cryo-FIB) milling before tilt-series acquisition on the resulting lamellae. In the reconstructed tomograms, paclitaxel-treated cells frequently contained large, dense bundles of parallel microtubules and vimentin filaments (Figure 1F and G; Supplementary Figure 1C) which were absent from control cells (Supplementary Figure 1C). These bundles could be observed to closely associate with the NE (Figure 1F and G; Supplementary Figure 1C).

Overall, these data indicate large-scale reorganisation of the cytoskeleton, beyond just microtubule filaments, in the perinuclear region that could lead to changes in nuclear-cytoskeletal coupling following paclitaxel treatment.

### Paclitaxel-induced nuclear deformation is dependent on Lamin A/C expression levels

LINC complexes connect the cytoskeleton to the NE and underlying lamina^28^. We hypothesised that the observed cytoskeletal reorganisation would affect this mechanical coupling and force distribution to the nucleus, resulting in nuclear deformation. To test this, high-content live cell imaging was performed over 48 h in the presence of 0, 1, 5, or 10 nM paclitaxel to measure nuclear solidity of nuclei in wild-type fibroblasts (Figure 2A and B; Supplementary Figure 2A and B). Nuclear solidity provided a measure of nuclear deformation^46^ (Supplementary Figure 2A) on a scale of 0 (most deformed) to 1 (least deformed), with Hoechst used as a nuclear marker. As expected, nuclear solidity decreased over time in paclitaxel-treated cells, and this nuclear deformation was concentration dependent (Figure 2A and B). Decreased nuclear solidity upon paclitaxel treatment was also observed in slow dividing, serum-starved cells, thus suggesting that the observed nuclear deformation occurs independently of cell division (Figure 2C; Supplementary Figure 2C and D).

**Figure 2.**
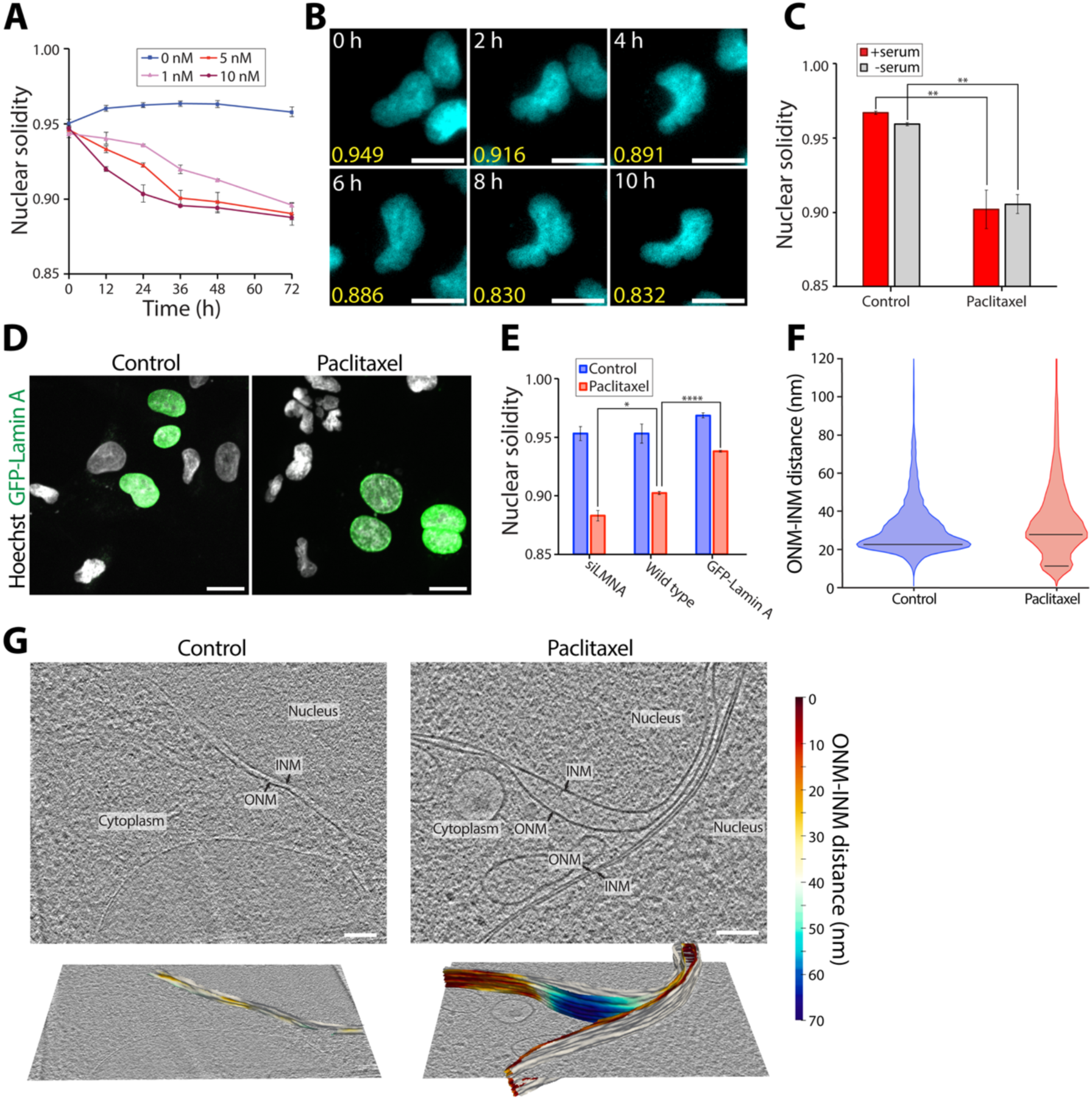
Paclitaxel results in nuclear deformation. **(A)** Nuclear solidity of Hoechst-stained nuclei over 72 h in 0, 1, 5, or 10 nM paclitaxel. Error bars show the SEM from three biological repeats, each with more than 50 cells. **(B)** Example frames from a live-cell time-series used for the nuclear solidity analysis in (A), showing a Hoechst-stained nuclei (cyan) and respective nuclear solidity measurements (yellow) taken every two hours between 0 and 10 h after the addition of 5 nM paclitaxel. **(C)** Bar chart comparing the nuclear solidity of cells cultured with complete medium (+serum) or serum-starved medium (-serum) in control conditions or after 16 h in 5 nM paclitaxel. Error bars show the SEM from three biological repeats (n=3), each with at least 50 cells. Statistical analysis: t-test control versus paclitaxel; +serum, P=0.0075 (**); -serum, P=0.0011 (**). **(D)** Confocal micrographs of cells transiently transfected with GFP-LMNA to overexpress GFP-Lamin A (green), fixed after 16 h incubation in control media or 5 nM paclitaxel. Cells were stained for DNA using Hoechst (grey). Scale bars = 20 μm. **(E)** Nuclear solidity of Hoechst-stained nuclei from wild-type cells, cells with Lamin A/C knocked down (siLMNA), or cells with GFP-Lamin A overexpressed, following 16 h incubation in control media or 5 nM paclitaxel. For GFP-Lamin A overexpression samples, only cells expressing GFP-Lamin A were analysed. Error bars show SEM from three biological repeats (n=3), each with at least 30 cells. Statistical analysis: t-test; wild type paclitaxel versus siLMNA paclitaxel, P=0.0146 (*); wild type paclitaxel versus GFP-Lamin A paclitaxel, P=1.74 × 10^−5^ (****). **(F)** Violin plot comparing the ONM-INM distance between control and paclitaxel-treated cells, quantified from nuclear membrane segmentations from high-resolution tomograms using surface morphometrics^49^. A total NE area of 1.64 μm^2^ over 23 tomograms from 6 cells was analysed for the control, and 3.97 μm^2^ over 41 tomograms from 9 cells was analysed for paclitaxel-treated cells. The black lines represent the modal values (21.0 nm for control; 10.5 nm and 25.5 nm for paclitaxel). **(G)** Example 2D slices of reconstructed tomograms of the NE in control and paclitaxel-treated cells. Segmentations of the ONM and INM that were used for the morphometrics analysis in (F) are shown in the lower panels, with the ONM coloured using a heatmap of the ONM-INM distance.

Since A-type lamins are major determinants of nuclear rigidity and the ability of the nucleus to withstand mechanical forces^47^, we next tested how Lamin A/C expression levels affect paclitaxel-induced nuclear deformation. Nuclear solidity was compared between wild-type cells, cells transiently overexpressing GFP-Lamin A, and cells with Lamin A/C depleted using siRNA (Figure 2D and E; Supplementary Figure 2E). In the absence of paclitaxel, nuclear solidity remained high across all three Lamin A/C expression levels. However, in the presence of paclitaxel, nuclear deformation was significantly greater in Lamin A/C depleted cells and significantly reduced when Lamin A was overexpressed (Figure 2E). Overall, this indicates that paclitaxel-induced nuclear deformation depends on Lamin A/C expression levels.

In addition to misshapen nuclei, cells containing multiple smaller nuclei, termed multimicronuclei, were observed following paclitaxel treatment (Supplementary Figure 2F). Live cell imaging showed that in every instance they arose immediately following cell division, but we never observed this during interphase (Supplementary Figure 2G). Importantly, quantifying the proportion of cells undergoing mitosis following 30 h incubation in 5 nM paclitaxel using immunofluorescence analysis showed that the proportion of mitotic cells in paclitaxel was 6.9 ± 0.4%, only slightly higher than that in control conditions (4.4 ± 0.6%) (Supplementary Figure 2H). This suggests that at clinically relevant concentrations, paclitaxel did not induce sustained mitotic arrest, and cells were instead able to exit mitosis. However, immunofluorescence microscopy analysis of paclitaxel-treated cells revealed that this mitosis was defective, since dividing cells contained disorganised mitotic spindles that were frequently multipolar, in contrast to bipolar spindles in untreated cells (Supplementary Figure 2I). Overall, our data indicates that nuclear deformation in paclitaxel occurs both during interphase and following defective mitosis.

The appearance of multimicronuclei in ovarian cancer cells was previously observed to depend on decreased Lamin A/C expression levels^14^. In line with this, we found that the frequency of multimicronucleated cells after 24 h in 5 nM paclitaxel was significantly lower when Lamin A was overexpressed and significantly higher when Lamin A/C was knocked down compared to the wild type (Supplementary Figure 2J).

To investigate changes to nuclear ultrastructure caused by paclitaxel, we used cryo-ET. Tomographic data of the NE showed altered intermembrane spacing between the inner and outer nuclear membranes (INM and ONM, respectively) following paclitaxel treatment (Figure 2F and G; Supplementary Figure 2K). In non-treated cells, this distance is uniformly around 25 nm^48^, but in paclitaxel-treated cells we frequently observed NE areas where the membranes appeared to have large separation or to the contrary were in very close proximity. To quantify this, we used morphometrics analysis^49^ of the NE segmented from tomograms of control and paclitaxel-treated cells (Figure 2F and G; Supplementary Figure 2K). In control cells, 67% of the NE had spacing of 20-40 nm, whereas in paclitaxel-treated cells this was decreased to 49% (Figure 2F). This was due to the significantly greater variability in the ONM-INM distance in paclitaxel-treated cells (Levene test: P<0.0001) because of the increased frequency of loci where the membranes were in very close proximity or highly separated, as seen in Figure 2G and Supplementary Figure 2K. Indeed, the ONM-INM distance showed a broader and bimodal distribution in paclitaxel-treated cells, with a peak within the expected range at 25.5 nm and an additional peak at 10.5 nm where the two membranes were in very close proximity (Figure 2F). This contrasted with the narrow distribution around the single peak at 21.0 nm for the control (Figure 2F). Together, this indicates that perturbation to the cytoskeleton due to paclitaxel treatment results in deformation to the overall nuclear shape and to the NE, resulting in altered NE spacing.

### Paclitaxel treatment disrupts the nuclear lamina via the LINC complex protein SUN2

Next, we investigated whether the nuclear lamina is also impacted by paclitaxel treatment. Immunofluorescence analysis of Lamin A/C and Lamin B1 showed irregular, patchy nuclear distribution of both proteins in paclitaxel-treated cells, compared to control cells (Figure 3A; Supplementary Figure 3A). Interestingly, Western blotting revealed that total cellular levels of Lamin A/C were decreased by paclitaxel in a concentration-dependent manner (Figure 3B and C) but, in contrast, Lamin B1 levels were not significantly affected (Figure 3B and C).

**Figure 3.**
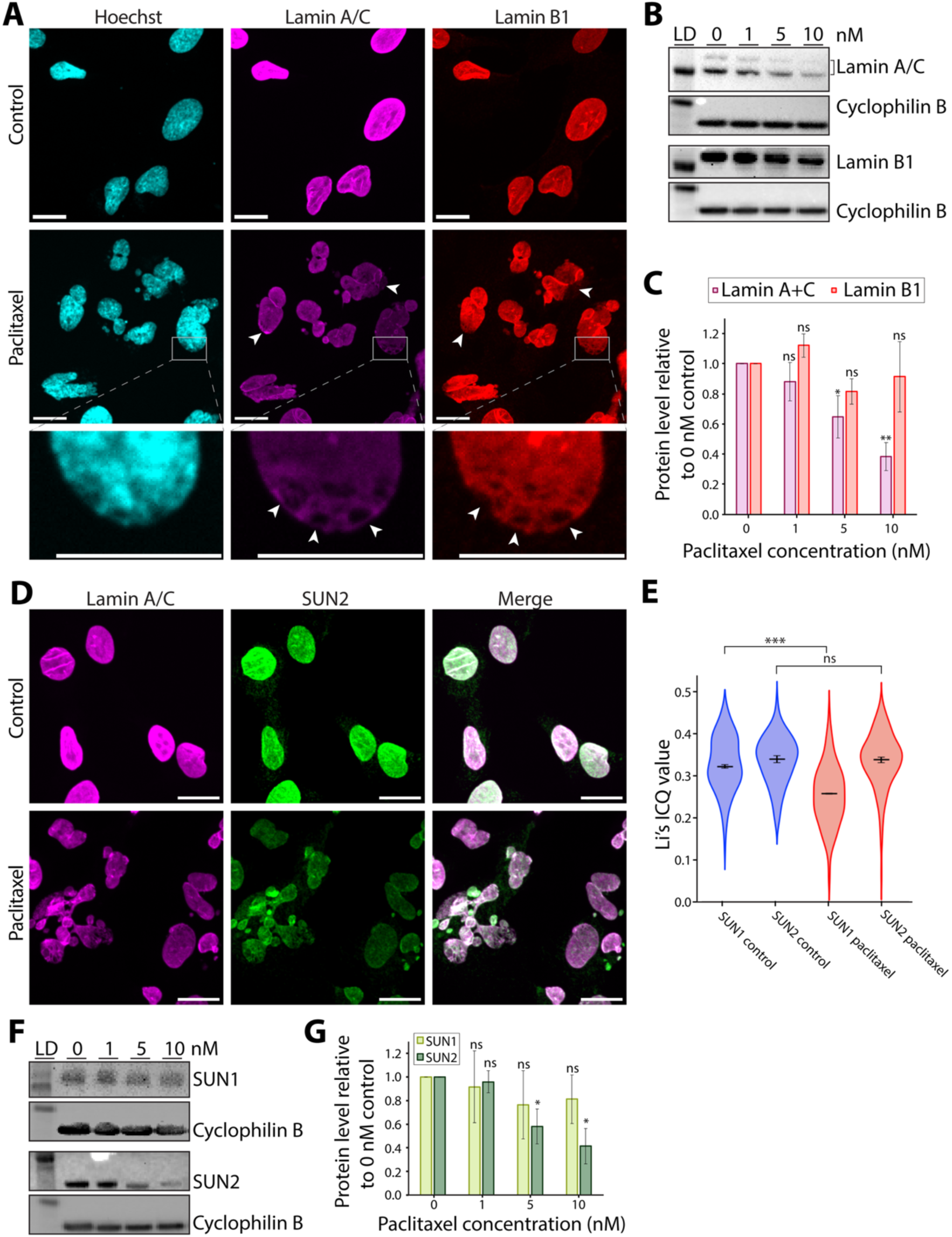
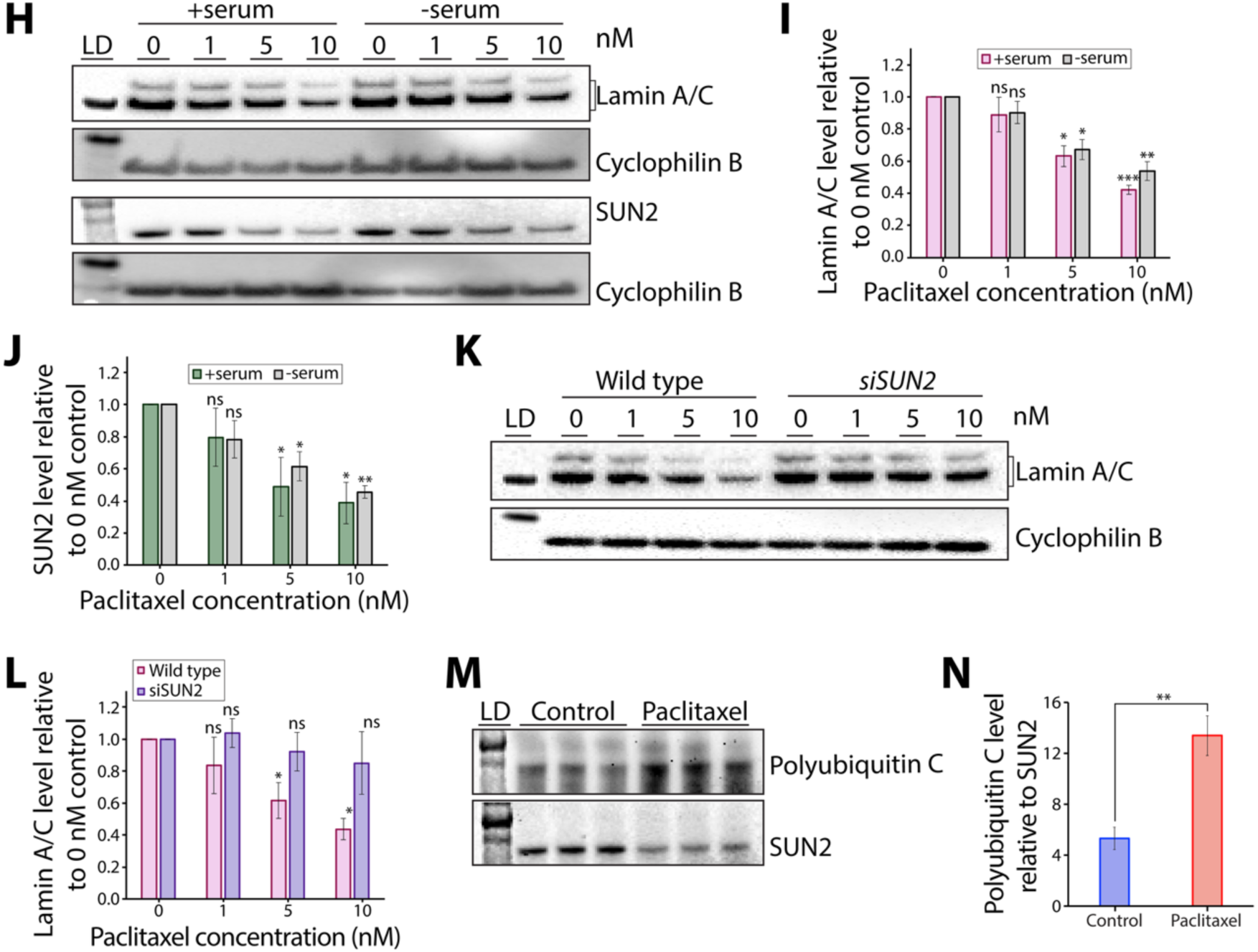
Paclitaxel treatment results in aberrant organisation of the nuclear lamina and decreased Lamin A/C levels via SUN2. **(A)** Confocal micrographs of cells fixed after 16 h incubation in control media or 5 nM paclitaxel. DNA was stained using Hoechst (cyan), and Lamin A/C (magenta) and Lamin B1 (red) were stained using immunofluorescence. Areas of the lamina that are patchy or have Lamin A/C or B1 missing are marked with arrowheads. Lower panels show magnified views of a patchy area of the lamina. Scale bars = 20 μm. **(B)** Western blots for Lamin A/C and Lamin B1 of whole cell lysates following 16 h incubation in media containing 0, 1, 5, or 10 nM paclitaxel. Cyclophilin B was used as a loading control. Lane 1 shows protein ladder (LD; 62 kDa for Lamin A/C and Lamin B1 panels, and 28 kDa for Cyclophilin B panels). **(C)** Quantification of Lamin A/C and Lamin B1 protein levels from (B). Each band was normalised to the corresponding Cyclophilin B loading control. Error bars represent the standard deviation from three biological repeats (n=3). Statistical analysis: one-sample t-test with null hypothesis mean = 1. Lamin A/C: 1 nM P=0.2469; 5 nM P=0.0489; 10 nM P=0.0077. Lamin B1: 1 nM P=0.0762; 5 nM P=0.0854; 10 nM P=0.2345. **(D)** Confocal micrographs of cells fixed after 16 h incubation in control media or 5 nM paclitaxel. Cells were stained for Lamin A/C (magenta) and SUN2 (green) using immunofluorescence. Scale bars = 20 μm. **(E)** Analysis of the co-localisation between Lamin A/C and SUN1 or SUN2. Fluorescence colocalization was quantified using Li’s ICQ value^55^, where more positive values represent better positive colocalization. Error bars represent the SEM from 3 biological repeats (n=3), each with more than 30 cells. Statistical analysis: t-test control versus paclitaxel: SUN1 P=0.0002; SUN2 P=0.8788. **(F)** Western blots for SUN1 and SUN2 of whole cell lysates following 16 h incubation in media containing 0, 1, 5, or 10 nM paclitaxel. Cyclophilin B was used as a loading control. Lane 1 shows protein ladder (LD; 98 kDa for SUN1 and SUN2 panels, and 28 kDa for Cyclophilin B panels). **(G)** Quantification of SUN1 and SUN2 protein levels from (F). Each band was normalised to the corresponding Cyclophilin B loading control. Error bars represent the standard deviation from three biological repeats (n=3). Statistical analysis: one-sample t-test with null hypothesis mean = 1. SUN1: 1 nM P=0.6816; 5 nM P=0.2948; 10 nM P= 0.2548. SUN2: 1 nM P=0.5320; 5 nM P=0.0395; 10 nM P= 0.0214. **(H)** Western blots for Lamin A/C and SUN2 of whole cell lysates from cells cultured in complete medium (+serum) or serum-starved medium (-serum) and following 16 h incubation in 0, 1, 5, or 10 nM paclitaxel. Cyclophilin B was used as a loading control. Lane 1 shows protein ladder (LD; 62 kDa for Lamin A/C panel, 98 kDa for SUN2 panel, and 28 kDa for Cyclophilin B panel). **(I)** Quantification of Lamin A/C protein levels from (H). Each band was normalised to the corresponding Cyclophilin B loading control. Error bars represent the standard deviation from three biological repeats (n=3). Statistical analysis: one-sample t-test with null hypothesis mean = 1. +Serum: 1 nM P=0.221; 5 nM P=0.0100; 10 nM P=0.0007. -Serum: 1 nM P=0.1306; 5 nM P=0.0126; 10 nM P=0.0056. **(J)** Quantification of SUN2 protein levels from (H). Each band was normalised to the corresponding Cyclophilin B loading control. Error bars represent the standard deviation from three biological repeats (n=3). Statistical analysis: one-sample t-test with null hypothesis mean = 1. +Serum: 1 nM P=0.1904; 5 nM P=0.0404; 10 nM P=0.0146. - Serum: 1 nM P=0.0849; 5 nM P=0.0182; 10 nM P=0.0015. **(K)** Western blots for Lamin A/C of whole cell lysates from wild-type or SUN2-knockdown (siSUN2) cells following 16 h incubation in 0, 1, 5, or 10 nM paclitaxel. Cyclophilin B was used as a loading control. Lane 1 shows protein ladder (LD; 62 kDa for Lamin A/C panel, and 28 kDa for Cyclophilin B panel). **(L)** Quantification of Lamin A/C protein levels from (K). Each band was normalised to the corresponding Cyclophilin B loading control. Error bars represent the standard deviation from three biological repeats (n=3). Statistical analysis: one-sample t-test with null hypothesis mean = 1. Wild type: 1 nM P=0.2455; 5 nM P=0.0271; 10 nM P=0.0046. siSUN2: 1 nM P=0.5481; 5 nM P=0.3769; 10 nM P=0.3137. **(M)** Top panel: Western blot for Polyubiquitin C following pull-down of SUN2 from whole cell lysates of control cells or cells treated with 5 nM paclitaxel for 16 h. Three biological repeats were used for each condition (n=3). Bottom panel: to control for SUN2 protein levels, the same membrane was stripped and blotted for SUN2. **(N)** Quantification of Polyubiquitin C levels from (M), with each band normalised to SUN2. Error bars represent the standard deviation from three biological repeats (n=3). Statistical analysis: t-test control versus paclitaxel, P=0.0014. Ns (non-significant) = P>0.05; * P<0.05; ** P<0.01; *** P<0.001; **** P<0.0001.

Reverse transcription quantitative PCR (RT-qPCR) showed that the paclitaxel-induced decrease in Lamin A/C levels did not occur at the mRNA level (Supplementary Figure 3B), indicating that regulation occurs at the protein level instead. Lamin A/C can be phosphorylated during interphase at several sites^50^, and this can alter its assembly into the lamina network and overall stability^51^. However, immunoblotting of whole-cell lysates against phospho-Ser404, a major site of Lamin A/C phosphorylation during interphase^50^, showed no changes in the phosphorylation levels after paclitaxel treatment (Supplementary Figure 3C and D). Phos-tag gel analysis corroborated these findings, revealing no changes in the overall phosphorylation state of Lamin A/C (Supplementary Figure 3E and F).

We also tested the acetylation and ubiquitination states of Lamin A/C, as these are known to affect its protein stability and assembly^52,53^. However, immunoprecipitation experiments against Lamin A/C followed by Western blotting for acetyl-lysine and polyubiquitin showed no significant change in Lamin A/C’s acetylation or ubiquitination state in paclitaxel-treated cells (Supplementary Figure 3G-I). This suggests an alternative mechanism for the observed reduction in Lamin A/C levels following paclitaxel treatment.

We hypothesised that perturbation to Lamin A/C in paclitaxel could instead occur through its direct connection to the cytoskeleton mediated via LINC complexes, in particular via SUN-proteins^28,30^. To test this, we used confocal microscopy to analyse the co-localisation of SUN1 and SUN2 with Lamin A/C (Figure 3D and E; Supplementary Figure 3J and K). Our data show that following paclitaxel treatment, both SUN1 and SUN2 become unevenly distributed at the NE, resulting in a patchy appearance (Figure 3D; Supplementary Figure 3J). However, while SUN2 retained high co-localisation with Lamin A/C, SUN1 showed decreased co-localisation in paclitaxel-treated cells (Figure 3D and E; Supplementary Figure 3J and K). Furthermore, Western blotting revealed that SUN2 protein levels were decreased in a concentration-dependent manner (Figure 3F and G), whereas SUN1 showed no significant reduction in total protein levels (Figure 3F and G). Since paclitaxel’s effects on SUN2 parallel those of Lamin A/C, this suggests that there is a close interplay between these two proteins in paclitaxel.

Western blot analysis of serum-starved cells showed a similar decrease in Lamin A/C and SUN2 levels following paclitaxel treatment (Figure 3H-J), indicating that this perturbation is not strictly dependent on cell division. Therefore, we hypothesised that the paclitaxel-induced changes to Lamin A/C protein levels occur through its direct connection to the cytoskeleton via SUN2-containing LINC complexes. Indeed, in contrast to the decrease of Lamin A/C levels in paclitaxel-treated wild-type cells, Lamin A/C levels remained high across all paclitaxel concentrations tested in SUN2 knockdown cells (Figure 3K and L; Supplementary Figure 3L). In the absence of SUN2, transmission of forces from the cytoskeleton to the nuclear lamina could be impaired and therefore less able to perturb Lamin A/C levels. However, when Lamin A/C is knocked down, a concentration-dependent decrease in SUN2 protein levels is still observed upon paclitaxel treatment (Supplementary Figure 3M and N). Altogether, our results indicate that SUN2-containing LINC complexes are essential in sensing and transmitting cytoskeletal forces to the nuclear lamina in paclitaxel.

RT-qPCR showed that the decrease in SUN2 levels following paclitaxel treatment did not occur at the mRNA level (Supplementary Figure 3O). Mechanical regulation of SUN2 is mediated by ubiquitination by the E3 ubiquitin ligase SCF^βTrCP 54^. To test whether the paclitaxel-induced decrease in SUN2 levels is mediated by SUN2 ubiquitination, we first performed we pulled-down SUN2 from whole-cell lysates of control or paclitaxel treated cells and performed Western blotting for Polyubiquitin C (Figure 3M and N). This showed that there was a significant increase in SUN2 ubiquitination following paclitaxel treatment (Figure 3M and N).

Finally, we confirmed that these results apply to cancer cells. As expected, paclitaxel treatment also resulted in a significant decrease in nuclear solidity (Supplementary Figure 3P), a patchy nuclear lamina (Supplementary Figure 3Q), and decreased Lamin A/C and SUN2 levels in breast cancer MDA-MB-231 cells (Supplementary Figure 3R-T).

### Nuclear deformation and a patchy nuclear lamina are specific to microtubule stabilisation

To test if the nuclear deformation and nuclear lamina disruption observed in interphase cells after paclitaxel treatment occur specifically due to microtubule bundling, we overexpressed Tau, a neuronal microtubule-associated protein known to induce microtubule stabilisation and bundling^56^. Cells transiently overexpressing GFP-Tau showed the presence of large microtubule bundles similar to paclitaxel-treated cells (Figure 4A). However, unlike paclitaxel treatment which induced microtubule bundling into rings around the nucleus (Figure 1A), the microtubule bundles in cells overexpressing GFP-Tau were distributed throughout the cytoplasm (Figure 4A). Interestingly, in cells where Tau-induced microtubule bundles were in close proximity to the nucleus, we observed substantial nuclear shape deformation (Figure 4A) and a significant decrease in nuclear solidity (Figure 4B). Furthermore, GFP-Tau overexpression also resulted in an uneven distribution of both Lamin A/C and Lamin B1 (Figure 4C), confirming that perturbation to the nuclear lamina during interphase is caused by microtubule stabilisation.

**Figure 4.**
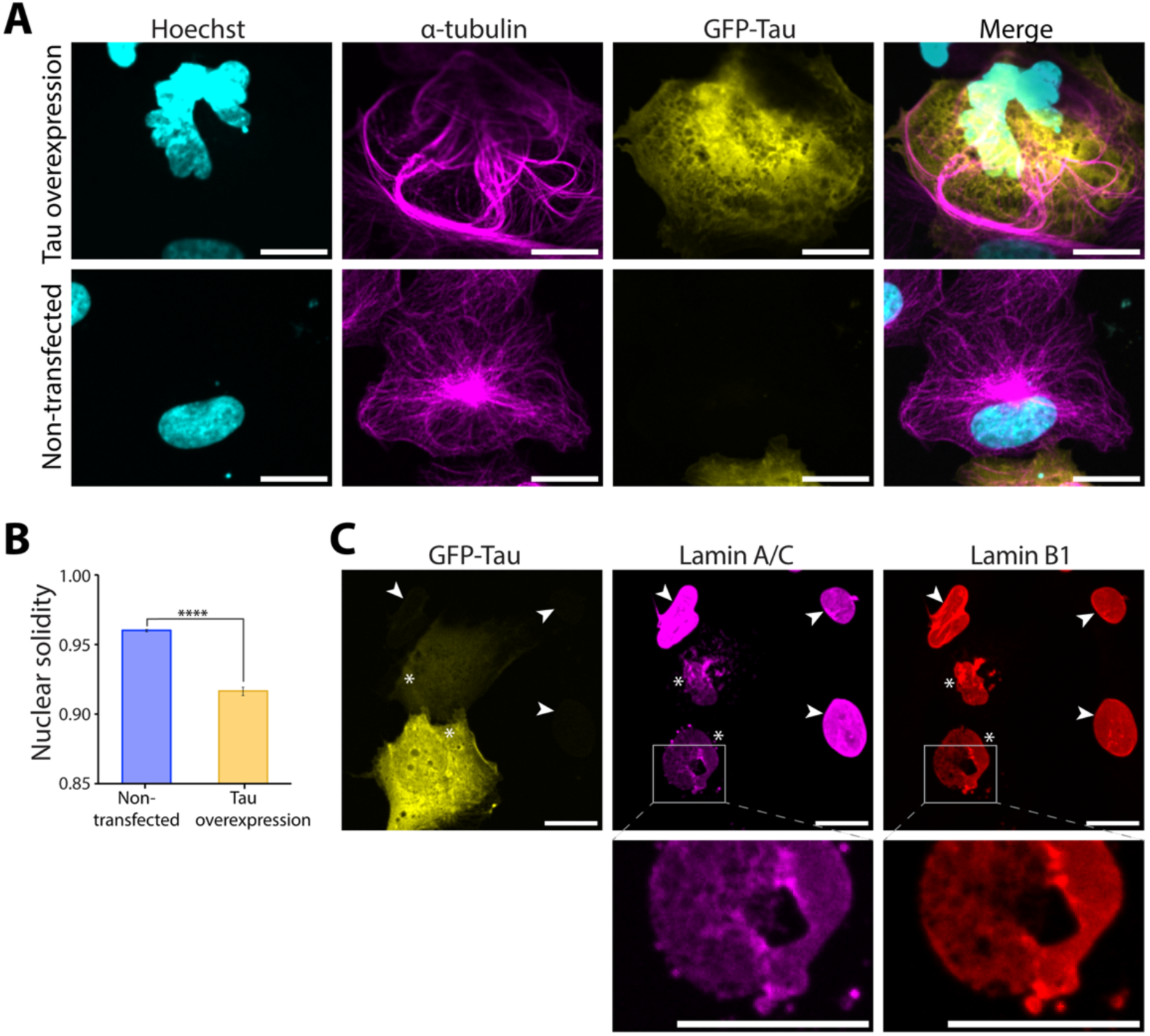
Microtubule stabilisation by GFP-Tau overexpression results in a patchy nuclear lamina and decreased nuclear solidity. **(A)** Confocal micrographs of cells fixed 48 h after transient transfection with GFP-Tau (yellow). DNA was stained using Hoechst (cyan), and microtubules using α-tubulin immunofluorescence (magenta). The top panel shows a cell overexpressing GFP-Tau, while the bottom panel shows a non-transfected control cell. Scale bars = 20 μm. **(B)** Confocal micrographs of cells fixed 48 h after transient transfection with GFP-Tau (yellow). Lamin A/C (magenta) and Lamin B1 (red) were stained using immunofluorescence. Cells overexpressing GFP-Tau are marked with asterisks, while non-transfected control cells are marked with arrowheads. Lower panels show magnified views of a patchy area of the lamina. Scale bars = 20 μm. **(C)** Bar chart comparing the nuclear solidity of non-transfected cells and cells overexpressing GFP-Tau. Error bars show the SEM from four biological repeats (n=4), each with at least 60 cells. Statistical analysis: t-test, P=7.68 × 10^−6^ (****).

### Lamin A/C expression levels determine sensitivity to paclitaxel treatment

As Lamin A/C expression levels determine nuclear deformation in paclitaxel, we then tested how this relates to overall cellular sensitivity to paclitaxel. We performed high-content screening to quantify cell confluency of the wild type, Lamin A/C knockdown, and GFP-Lamin A overexpression over 48 h in 0, 1, 5, or 10 nM paclitaxel (Figure 5A). Without paclitaxel, cell growth was similar across all three Lamin A/C expression levels tested. In 1 and 5 nM paclitaxel, cell growth was reduced, but to a significantly lesser extent when Lamin A was overexpressed compared to the wild type and Lamin A/C knockdown (Figure 5A). In 10 nM paclitaxel, cell confluency remained low across all three Lamin A/C expression levels (Figure 5A) as the high concentration prevented almost any cell growth.

**Figure 5.**
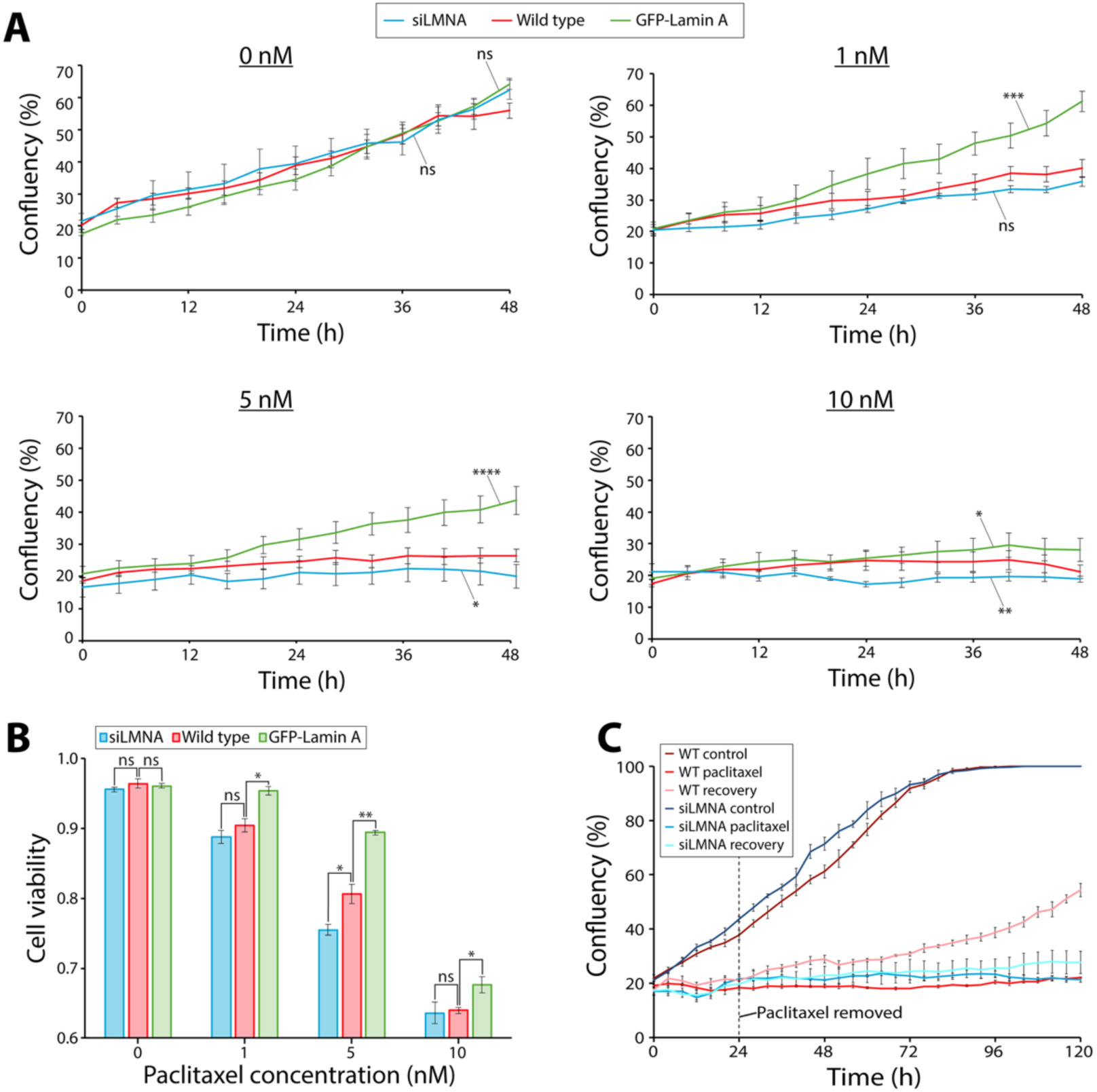
Lamin A/C expression level affects cell sensitivity to and recovery from paclitaxel. **(A)** Cell confluency of wild-type (red), GFP-Lamin A overexpression (green), and Lamin A/C knockdown (blue) cells over 48 h in 0, 1, 5, or 10 nM paclitaxel. Confluency was calculated from brightfield high-content live cell images. Error bars represent the SEM from five biological repeats. Statistical tests: one-way three-factor repeated measures ANOVA with Bonferroni post hoc test. 0 nM: GFP-Lamin A vs wild type P=0.3186; siLMNA vs wild type P=0.6074. 1 nM: GFP-Lamin A vs wild type P=0.0003; siLMNA vs wild type P=0.1662. 5 nM: GFP-Lamin A vs wild type P=4.5 × 10^−5^; siLMNA vs wild type P=0.0267. 10 nM: GFP-Lamin A vs wild type P=0.0207; siLMNA vs wild type P=0.0029. **(B)** Bar graph comparing cell viability between wild-type (red), GFP-Lamin A overexpression (green), and Lamin A/C knockdown (blue) cells following 48 h incubation in 0, 1, 5, or 10 nM paclitaxel. Cell viability was calculated by subtracting the proportion of the total number of cells that were dead/apoptotic (identified by FLICA staining) from 1. Error bars represent SEM from three biological repeats (n=3), each with at least 95 cells. Statistical analysis: t-test versus wild type. 0 nM: siLMNA P= 0.3244; GFP-Lamin A P= 0.6742. 1 nM: siLMNA P= 0.2911; GFP-Lamin A P= 0.0107. 5 nM: siLMNA P= 0.0325; GFP-Lamin A P= 0.0036. 10 nM: siLMNA P= 0.8297; GFP-Lamin A P= 0.0428. **(C)** Cell confluency of wild-type and Lamin A/C knockdown cells in control media (control), media with the sustained presence of 5 nM paclitaxel (paclitaxel), or media from which paclitaxel was removed after 24 h (recovery). Ns (non-significant) = P>0.05; * = P<0.05, ** P<0.01, *** P<0.001, **** P<0.0001.

Next, we tested the effects of Lamin A/C expression levels on cell viability in paclitaxel (Figure 5B) by calculating the proportion of dead or apoptotic cells using the caspase-3/7 dye FLICA (Supplementary Figure 4A). Increasing concentrations of paclitaxel resulted in lower cell viability, and this was more pronounced in Lamin A/C depleted cells (Figure 5B). However, at high concentrations of paclitaxel (10 nM), Lamin A/C knockdown no longer had an observable impact on cell viability. Conversely, cells overexpressing Lamin A displayed higher cell viability in paclitaxel, even at high concentrations (Figure 5B). Overall, these data indicate that Lamin A/C expression levels affect sensitivity to paclitaxel in terms of both cell growth and cell death.

Finally, we tested recovery from paclitaxel treatment. Immunofluorescence confocal microscopy showed that microtubule bundling was reverted following paclitaxel removal (Supplementary Figure 4B). Importantly, the localisation of Lamin B1 and Lamin A/C (Supplementary Figure 4B) and the protein levels of Lamin A/C and SUN2 also recovered after removal of 5nM paclitaxel (Supplementary Figure 4C and D).

To assess the effect of Lamin A/C levels on paclitaxel recovery, we also measured cell confluency of wild-type and Lamin A/C knockdown cells following removal of paclitaxel (Figure 5C). Whereas wild-type cells expanded in cell number following paclitaxel removal, Lamin A/C knockdown cells showed little cell growth (Figure 5C). Altogether, this shows that Lamin A/C expression levels also affect recovery from paclitaxel, in addition to cell death and inhibition of cell growth.

## Discussion

Despite its widespread use, the mechanism by which paclitaxel selectively kills cancer cells is not yet fully understood. A better understanding of paclitaxel’s effects is therefore important for improving treatment and reducing drug resistance and toxicity. In this study, we investigated how paclitaxel disrupts nuclear-cytoskeletal coupling during interphase. We show that paclitaxel treatment results in nuclear aberrations, including NE deformation, changes to nuclear intermembrane spacing, multimicronucleation, and disruption to the nuclear lamina via SUN2-containing LINC complexes. Cells with lower Lamin A/C expression levels, typical of many different cancer cells^36–40^, showed increased sensitivity to paclitaxel in terms of cell growth, death, and recovery, indicating how disrupted nuclear-cytoskeletal coupling can lead to a selective targeting of cancer cells, even in slowly proliferating tumours. Furthermore, we directly link nuclear aberrations to microtubule bundling around the nucleus.

The anti-cancer activity of paclitaxel was originally thought to be solely due to sustained mitotic arrest caused by disruption of the mitotic spindle^7^. Evidence for this came from *in vitro* cell culture and xenograft models which rely on cells with extensive replicative capacity and short doubling times^7,57–59^. However, these cells may be more vulnerable to mitotic arrest than tumour cells in patients, which can have longer doubling times^15,60^. Furthermore, tumour biopsies from breast cancer patients treated with microtubule stabilising drugs show that mitotic cells are rare, despite patient response to therapy^15,61^. One alternative hypothesis to explain this lack of mitotic cells in paclitaxel-treated tumour samples is that at intratumoral concentrations, cancer cells undergoing mitosis can escape the mitotic checkpoint, and this mitotic slippage results in defective chromosome segregation and apoptosis of the resulting aneuploid cells^8,9^. Interestingly, we show here that mitotic slippage in clinically relevant concentrations of paclitaxel can also occur in healthy human cells since the mitotic index of healthy human fibroblasts remained low in paclitaxel and they could exit mitosis to result in multimicronucleation (Supplementary Figure 2F-H). However, the frequency of multimicronucleation, which always occurred following exit from mitosis (Supplementary Figure 2G), depended on Lamin A/C expression levels (Supplementary Figure 2J), as previously reported in ovarian cancer cells^14^. This is likely explained by the important role the nuclear lamina plays in the fragmentation and reassembly of nuclear membranes during mitosis^62,63^. NE reformation around aberrantly segregated chromosomes could therefore be affected by Lamin A/C expression levels, explaining the differences in multimicronucleation frequency and how cancer cells could be more susceptible due to their aberrant NEs that frequently show lower Lamin A/C expression levels^40^.

However, multimicronucleation alone cannot account for paclitaxel’s activity against tumours with slow duplication rates^10–12^, as it is unlikely that enough cells undergo mitosis in paclitaxel for this mechanism to be sufficient. This is supported by intravital imaging of murine tumours treated with docetaxel, another taxane closely related to paclitaxel, which demonstrated that apoptosis is induced in most tumour cells independently of aberrant mitosis, including both mitotic arrest and multimicronucleation^11^. For its activity against these cancers, paclitaxel must therefore have additional anti-cancer effects outside of mitosis^13–15^. In support of this, drugs that specifically block mitosis have not been successful in the clinic, indicating that the efficacy of microtubule-targeting drugs like paclitaxel comes from additional effects in interphase as well as mitosis^15^. Recent studies showing that paclitaxel affects the nuclear integrity of cancer cells in interphase indicate that this activity could be due to disruption of nuclear-cytoskeletal coupling^14,20,21^.

Previously, paclitaxel was shown to affect microtubule organisation in interphase cells in a cell-type specific manner, with some cells containing microtubule rings around the nucleus while others contain multiple microtubule asters^23^. Here, we show that paclitaxel promotes the formation of dense bundles of microtubules and vimentin filaments surrounding the interphase nucleus in human fibroblasts (Figure 1 and Supplementary Figure 1), which coincide with nuclear deformation (Figure 2A and B) and changes to NE intermembrane spacing (Figure 2F and G). These cytoskeletal rearrangements in fibroblasts mirror those in breast cancer MDA-MB-231 cells (Supplementary Figure 1B). Since paclitaxel-treated cells lack prominent actin stress fibres adjacent to the nucleus^22^ (Figure 1D), nuclear deformation likely occurs via the microtubule and vimentin cytoskeletons. Indeed, microtubules connect to LINC complexes in the NE via interactions between the microtubule motors kinesin and dynein and the N-terminus of KASH proteins^64,65^, while vimentin filaments connect to KASH proteins via plectin^66^. Furthermore, perinuclear assemblies of both microtubules and vimentin filaments have been previously observed to induce nuclear deformation in interphase^67,68^. The large, dense filament bundles in paclitaxel-treated cells may exert force on the nucleus either directly via steric effects or via their mechanical connection to LINC complexes^64–66^.

LINC complexes in turn connect to the nuclear lamina in the nucleoplasm^30,69^. Here, we show that paclitaxel treatment decreases Lamin A/C expression levels independently of cell division (Figure 3B, C, H, and I). In other systems, Lamin A/C levels respond to mechanical cues^70^, and this mechanosensing is thought to be achieved through post-translational modifications of Lamin A/C, including phosphorylation, acetylation, and ubiquitination^53,71,72^. In particular, Lamin A/C phosphorylation and acetylation regulate its assembly and stability in the nuclear lamina and its subsequent degradation^50,51,53^. Nevertheless, our results did not show any significant changes in phosphorylation, acetylation, or ubiquitination of Lamin A/C in paclitaxel (Supplementary Figure 3C-I). Instead, we postulate a different mechanism involving the direct connection of Lamin A/C to SUN2-containing LINC complexes (Figure 3K and L). The role of LINC complexes in paclitaxel-induced nuclear lamina perturbations is further supported by the fact that Lamin A/C but not Lamin B1 levels are decreased (Figure 3B and C). This is because, in contrast to robustly expressed Lamin B1 which is not mechanically linked to the cytoskeleton directly via LINC complexes^30,73^, Lamin A/C directly interacts with LINC complexes^30,69^, and its expression and turnover are known to be controlled by tissue stiffness and mechanical stress^70,73^.

A role for LINC complexes in paclitaxel-induced nuclear deformation is further supported by the uneven localisation and decreased protein levels of SUN2 (Figure 3D, F and G). This aberrant SUN2 localisation in the NE likely also accounts for the altered nuclear membrane spacing in paclitaxel (Figure 2F and G; Supplementary Figure 2K). Since SUN proteins in LINC complexes contain perinuclear coiled-coil domains of a set length, their presence in the NE controls the inter-membrane distance within narrow limits^69,74^. This control is lost when SUN proteins are absent^69^, enabling the two membranes to be more easily forced together or separated under cytoskeletal forces, as observed here (Figure 2F and G). SUN1 is still present in paclitaxel-treated cells, and this likely explains why a large proportion of the NE still has spacing within the expected range of 20-40 nm (Figure 2F). However, its expression levels remain the same and therefore SUN1 is unlikely to be able to completely compensate for the absence of SUN2, explaining the increased variability in NE spacing in paclitaxel-treated cells (Figure 2F).

It is still unclear why SUN2 and not SUN1 protein levels are specifically disrupted by paclitaxel treatment (Figure 3F and G). However, mechanical perturbations in the form of cyclic tensile strain have previously been shown to decrease levels of SUN2, but not SUN1, in human mesenchymal stem cells^75^. We show that this decrease in SUN2 levels occurs independently of Lamin A/C (Supplementary Figure 3M and N). Instead, paclitaxel treatment resulted in a significant increase in SUN2 ubiquitination (Figure 3M and N). This supports previous observations that the turnover of SUN2 via ubiquitination by the E3 ubiquitin ligase SCF^βTrCP^ is important in the control of nuclear mechanics, in particular nuclear shape^54^. SUN1 is not under the same control since it lacks the SCF^βTrCP^ recognition site present in SUN2^54^.

Although both SUN1 and SUN2 have been shown to interact directly with Lamin A/C^69^, upon paclitaxel treatment, SUN2 and Lamin A/C co-localised well whereas SUN1 did not (Figure 3D and E; Supplementary Figure 3J). This specific co-localisation indicates that SUN2 might be the main mediator of mechanical connections between Lamin A/C and microtubules and/or vimentin filaments in human fibroblasts. We propose that these mechanical connections are what drive perturbations to Lamin A/C in paclitaxel. Supporting this, the paclitaxel-induced decrease of Lamin A/C protein levels was no longer observed when SUN2 was knocked down (Figure 3K and L). This suggests that in the absence of SUN2-containing LINC complexes, aberrant cytoskeletal forces are no longer effectively transmitted to Lamin A/C to disrupt its protein levels at the nuclear lamina.

Furthermore, microtubule bundling induced by GFP-Tau overexpression resulted in similar perturbations to both Lamin A/C localisation and overall nuclear shape when these bundles were localised near the nucleus (Figure 4). This confirms that these effects of paclitaxel are due to its activity on the microtubule cytoskeleton in interphase, especially because GFP-Tau overexpression did not lead to multimicronucleation or appear to affect mitosis. Furthermore, this agrees with previous studies in neuroblastoma cells showing that Tau overexpression results in nuclear deformation via microtubule bundling^56^.

The disruption to the nuclear lamina and NE observed in paclitaxel, caused both by aberrant nuclear-cytoskeletal coupling during interphase and following exit from defective mitosis, likely compromises the ability of the nucleus to buffer cytoskeletal forces, resulting in loss of nuclear integrity, widespread DNA damage, and, if severe enough, cell death^26,76–78^. Consequently, cells already containing a defective nuclear lamina or NE may be more susceptible to these effects. Indeed, we showed that Lamin A/C knockdown increases the severity of nuclear deformation in paclitaxel (Figure 2D and E) as well as the frequency of multimicronucleation (Supplementary Figure 2J), and results in increased cell death, decreased cell growth, and decreased recovery from paclitaxel (Figure 5). Since nuclear abnormalities are common across most cancer types^35,40^, particularly alterations to the nuclear lamina and decreased Lamin A/C levels^36–40^, this study provides an explanation for the selective targeting of cancer cells by paclitaxel during interphase.

Overall, our work builds on previous studies investigating loss of nuclear integrity as an anti-cancer mechanism of paclitaxel separate from mitotic arrest^14,20,21^. We propose that cancer cells show increased sensitivity to nuclear deformation induced by aberrant nuclear-cytoskeletal coupling and multimicronucleation following mitotic slippage. Therefore, we conclude that paclitaxel functions in interphase as well as mitosis, elucidating how slowly growing tumours are targeted^10–12^. Since better understanding the mechanisms of paclitaxel and its effects on nuclear architecture is important for its safe and efficacious use as an anti-cancer drug, our study might inform patient stratification and the development of safer drug formulations. Similarly, our findings may be relevant for the search and/or design of other drugs that affect nuclear-cytoskeletal coupling.

## Materials and methods

### Cell culture and paclitaxel treatment

Human skin fibroblasts derived from AG10803 and immortalised with SV40LT and TERT were a gift from Dr Delphine Larrieu. These, along with MDA-MB-231 human breast cancer cells (Merck), were cultured at 37°C and 5% CO_2_ in complete medium consisting of Dulbecco’s modified Eagle’s medium (DMEM) with GlutaMAX (Gibco), supplemented with 10% fetal bovine serum (FBS) and 1% penicillin/streptomycin (Gibco).

For serum starvation, cells were resuspended in PBS before being pelleted at 300 x g for 5 min at room temperature. Cells were then washed in PBS twice, before being plated in low-serum medium (minimal essential medium [MEM] with 0.5% FBS and 1% penicillin/streptomycin). Cells were incubated in low-serum medium for three days prior to further experiments.

For paclitaxel treatment, paclitaxel (Merck PHL89806) resuspended in dimethyl sulfoxide (DMSO) to 10 mM was diluted in complete DMEM to final concentrations of 1, 5, or 10 nM as indicated.

### Transfections, RNA interference, and cell sorting

To overexpress Lamin A, cells were transfected using Lipofectamine 3000 (Invitrogen) with pBABE-puro-GFP-lamin A plasmid which was a gift from Tom Misteli (Addgene plasmid 17662)^79^. To overexpress Tau, cells were electroporated using a 4D nucleofector X kit (Lonza) with pRK5 GFP-Tau plasmid which was a gift from George Bloom (Addgene plasmid 187023)^80^. Overexpression was confirmed by fluorescence microscopy (Figure 2D; Figure 4A).

For knockdowns, cells were transfected with the indicated small interfering RNA (siRNA) at a final concentration of 100 nM using lipofectamine 3000 (Supplementary Table 1). Knockdown was confirmed using Western blot analysis (Supplementary Figure 2E; Supplementary Figure 3L). Drug treatments were started 48 hours after transfection.

### High-content live cell imaging

Cells adhered to 24-well plates (Corning) were stained for 10 min with Hoechst 33342 (ThermoFisher). The indicated concentration of paclitaxel was then added, and 24-well plates were transferred to CELLCYTE X Live Cell Imager (Cytena) fitted with a 10X objective lens (air, NA 0.3) held at 37°C and 5% CO_2_ and imaged every 2 h. For paclitaxel recovery, wells were washed and the media replaced with complete medium after 24 h.

To quantify nuclear shape deformation, images were transferred to ImageJ for analysis and Hoechst-stained nuclei were selected by threshold and the ‘analyse particles’ function used to calculate nuclear solidity. Nuclei that were overlapping with those from adjacent cells or that were within dead cells, identified by their distinctive cell shape, were excluded from the analysis.

Cell growth was quantified by measuring confluency from brightfield images using the automated image analysis tool in CELLCYTE X (Cytena).

### Cell staining and immunofluorescence

Following transfection and/or paclitaxel treatment, cells adhered on #1.5 glass coverslips were stained with Hoechst 33342 in complete medium for 10 min at 37°C. Cells were then washed with PBS, fixed with 4% paraformaldehyde for 15 min at room temperature, then washed with PBS before autofluorescence was quenched using 50 mM ammonium chloride. Cells were then permeabilised and blocked in permeabilization buffer (0.1% Triton X-100 and 2% bovine serum albumin in PBS) for 30 min then washed in PBS. Cells were then incubated at room temperature for 1 h in permeabilization buffer with the indicated primary antibodies (Supplementary Table 2) or Alexa488 phalloidin (Invitrogen). After washing with PBS, cells were incubated at room temperature for 1 h in secondary antibodies in permeabilization buffer (Supplementary Table 2) then washed with PBS. For STORM samples, cells were then incubated with 4% paraformaldehyde for 10 min then washed and stored in PBS. Otherwise, cells were mounted using Prolong Gold (ThermoFisher).

For quantification of cell viability, cells adhered on #1.5 glass coverslips pre-stained with Hoechst 33342 were then stained for active caspases 3 and 7 using FLICA reagent from Image-iT LIVE Red Caspase Detection Kit (Invitrogen). Cells were then washed twice, fixed with 4% paraformaldehyde for 15 min at room temperature, washed with PBS, then mounted. Following confocal microscopy, the proportion of dead or apoptotic cells was calculated by measuring the proportion of cells that were stained using FLICA.

### Light microscopy

Immunofluorescence imaging was performed using NIS-Elements software and a Nikon X1 Spinning Disk inverted microscope equipped with a 40X objective lens (oil immersion, numerical aperture [NA] 1.3) and sCMOS camera.

For Stochastic Optical Reconstruction Microscopy (STORM), immunofluorescence samples were immersed in STORM buffer (10% glucose, 10 mM sodium chloride, 50 mM Tris-HCl pH 8.0, 5.6 % glucose oxidase, 3.4 mg/ml catalase, 0.1% 2-mercaptoethanol) and imaged using a Nanoimager system (ONI) equipped with a 100X TIRF objective lens (oil immersion, NA 1.49). For imaging, Highly Inclined and Laminated Optical Sheet (HILO) illumination was used with laser power 60-150 mW, and 15,000 frames were acquired. CODI software (ONI) was used for image processing, including drift correction and HDBSCAN clustering. For α-tubulin cluster analysis, HDBSCAN clusters were constrained to have a minimum number of 15 localisations, maximum circularity of 0.5, and minimum length of 1000 nm so that the clusters mapped to individual filament bundles.

### Pull-downs and Western blotting

For Western blotting, cells were collected by treatment with 0.25% trypsin-EDTA for 3 min then dilution in PBS followed by centrifugation at 400 x g for 10 min. Cells were lysed with 1X NUPAGE LDS sample buffer supplemented with 50 mM dithiothreitol (DTT). After boiling at 95°C for 10 min, proteins were separated using NUPAGE 4-12% Bis-Tris gels (ThermoFisher) in MES SDS buffer alongside a protein ladder (Invitrogen SeeBlue Plus2). Proteins were transferred to PVDF membranes (Bio-Rad) which were then incubated in blocking buffer (5% milk power and 0.1% Tween 20 in Tris-buffered saline [TBS]). The membranes were then incubated in primary antibodies in blocking buffer (Supplementary Table 2), then washed with washing buffer (1% milk and 0.1% Tween 20 in TBS), then incubated with secondary antibodies in blocking buffer (Supplementary Table 2), then washed in washing buffer again. The membranes were then activated using ECL (Cytiva) then imaged using a Bio-Rad Chemidoc MP. Relative protein levels were quantified in ImageJ from three biological repeats, with normalisation to Cyclophilin B loading controls.

For Phos-tag gel analysis, cells were washed in PBS then lysed in cold lysis buffer (10 mM HEPES pH 7.5, 2 mM magnesium chloride, 25 mM potassium chloride, 0.1% NP-40, 0.1% Triton X-100, 0.1 mM DTT, 0.1 mM PMSF, and cOmplete EDTA-free inhibitor cocktail [Sigma-Aldrich]) with cell scraping. Following incubation on ice for 1 h, cell lysates were centrifuged at 14,000 x g for 15 min at 4°C, then the supernatant was boiled in 1X NUPAGE LDS sample buffer with 50 mM DTT before separation using 50 µM Phos-tag 10% acrylamide gels (FUJIFILM) in Tris-glycine SDS buffer. After running, the gel was washed twice in Tris-glycine SDS buffer with 1 mM EDTA, then once in Tris-glycine SDS buffer. Proteins were then transferred to PVDF membranes for Western blot analysis as above.

For pull-downs, cells were lysed as for the Phos-tag gel analysis. Cell lysates were centrifuged at 14,000 x g for 15 min at 4°C, before 250 µl of the lysate supernatant was pre-cleared by incubating with 25 µl protein A Dynabeads (Invitrogen) for 20 min at room temperature with end-to-end rotation. The beads were then magnetically separated and the pre-cleared lysate incubated with primary antibodies (Supplementary Table 2) overnight at 4°C with end-to-end rotation. The lysate-antibody mix was then added to 50 µl protein A Dynabeads and incubated for 1 h at room temperature with end-to-end rotation. The lysate was then removed by magnetic separation, and the beads washed three times in cold lysis buffer. The pulled-down proteins were then eluted by boiling the beads in 50 µl 1X NUPAGE LDS sample buffer for 5 min, before the beads were magnetically removed. Samples were then separated by gel electrophoresis for Western blotting as above. For loading controls, the membranes were then stripped by incubation in stripping buffer (15 mg/ml glycine, 1 mg/ml SDS, and 1% Tween 20 in water, adjusted to pH 2.2) for 10 min twice, before washing them twice in PBS for 10 min then twice in TBS with 0.1% Tween 20 for 5 min. The membranes were then blocked and probed with either anti-Lamin A/C or anti-SUN2 antibodies as above.

### Reverse transcription quantitative PCR

To extract mRNA, cells were washed with ice-cold PBS then lysed with 1 ml TRIzol (Invitrogen) for 2 min at room temperature, before shaking with 250 µl chloroform. After 5 min, the mixture was centrifuged at 10,000 x g. The top aqueous layer was then removed, added to 550 µl isopropanol, incubated at room temperature for 5 min, then centrifuged at 14,000 x g for 30 min. The tube was placed on ice and the isopropanol poured off, before adding 1 ml 75% ethanol. After centrifugation at 9,500 x g for 5 min, the ethanol was poured off before the RNA pellets were dried then resuspended in water. For reverse transcription, the TaqMan reverse transcriptase kit (Thermo Fisher Scientific) was used with random hexamer primers according to the manufacturer’s protocol. For qPCR, 0.6 µl cDNA was added to 3.4 µl water, 5 µl iTaq Universal SYBR Green Supermix (Bio-Rad), 0.5 µl 0.5 µM forward primers, and 0.5 µl 0.5 µM reverse primers (Supplementary Table 3). Three biological repeats were performed for the control and paclitaxel-treated samples. For each of these biological repeats, three technical replicates were performed using primers targeting the gene of interest, and three using primers targeting the housekeeping gene GAPDH. Forty cycles with denaturation at 95°C for 10 s and annealing/extension at 60°C for 60 s were run in a Bio-Rad CFX RT-qPCR machine. The cycle threshold (Ct) values were then used for 2^−ΔΔCt^ analysis, as previously described^81^.

### EM grid preparation

Quantifoil R 1/4 Au 200 mesh grids (Quantifoil Micro Tools GmbH) were glow discharged for 40 s at 30 mA with an Edwards S150B glow discharger then coated using 0.1 mg/ml poly-L-lysine for 30 min. Grids were then washed in PBS three times, then cells were seeded and allowed to adhere overnight, before 16 h treatment with 5 nM paclitaxel or control media. Grids were then transferred to a Leica GP2 plunger at 25°C and 95% humidity using glycerol (10% in PBS) as a cryo-protectant for 20 s. After 9 s blotting, grids were plunge frozen in liquid ethane.

### Cryo-focussed ion beam (FIB) milling and scanning electron microscopy (SEM)

To prepare lamellae for cryo-ET acquisition, cryo-FIB milling was performed using a Scios dual beam FIB/SEM (Thermo Fisher Scientific) equipped with a PP3010T cryo stage and loading system (Quorum Technologies) according to previously published protocols^82^. Briefly, the grids were sputter-coated with an inorganic platinum layer at 10 mA using the Quorum system sputter coater, then coated with an organometallic platinum layer (trimethyl [(1,2,3,4,5-ETA.)-1 methyl-2, 4-cyclopentadien-1-YL] platinum) using the Scios gas injection system. Milling was conducted at an angle of 10° relative to the grid in a stepwise manner with decreasing currents (0.5 nA, 0.3 nA, 0.1 nA, 30 pA), resulting in lamellae with a final width of 10-12 µm and thickness of 120-200 nm.

### Electron cryo-tomography and morphometrics analysis

Lamellae were imaged using a Titan Krios (TFS) equipped with a Gatan BioQuantum energy-filter and Gatan K3 direct electron detector. Tilt series were collected using SerialEM^83^ with PACE-tomo^84^ in a dose-symmetric scheme between ± 60° relative to the lamella in 3° increments. The total dose for each tilt series was ∼100 e/Å^2^, distributed evenly across each 12-frame tilt. The pixel size was 1.63 Å/pixel and defocus −3 to −5 µm.

For tomogram reconstruction, movies were imported into Warp^85^ for gain and motion correction, tilt selection, and CTF estimation. Tomograms were then reconstructed and deconvolved following tilt series alignment using AreTomo^86^.

Microtubules, vimentin filaments, and the inner and outer nuclear membranes were manually segmented in IMOD^87^. To analyse NE morphology, the NE segmentations were then converted to 3D voxel segmentations using the ‘imodmop’ program within IMOD. A surface morphometrics pipeline^49^ was then used to generate surface meshes of the inner and outer NEs from the 3D voxel segmentations and subsequently measure the distances between these meshes, giving the INM-ONM distance.

## Abbreviations

Cryo-FIB: cryo-focused ion beam
cryo-ET: electron cryo-tomography
DMEM: Dulbecco’s modified Eagle’s medium
DMSO: dimethyl sulfoxide
DTT: dithiothreitol
FBS: fetal bovine serum
hTERT: human telomerase reverse transcriptase
INM: inner nuclear membrane
LINC complex: Linker of Nucleoskeleton and Cytoskeleton complex
Li’s ICQ: Li’s Intensity Correlation Quotient
MEM: minimal essential medium
MTOC: microtubule organising centre
NE: nuclear envelope
ns: non-significant
ONM: outer nuclear membrane
RT-qPCR: reverse transcription quantitative PCR
SEM: standard error of the mean
STORM: stochastic optical reconstruction microscopy
TBS: Tris-buffered saline.

## Acknowledgements

We thank Dr David Barford, Dr Andrew Carter, and Prof Laura Machesky for in-depth discussions and critical feedback. We thank Dr Delphine Larrieu for providing hTERT human skin fibroblasts used in this study. We thank the EM Facility at the LMB, the LMB Scientific Computing facility and the LMB Light Microscopy Facility for technical support. We also acknowledge the UK national electron Bio-Imaging Centre (eBIC) for access to Aquilos-II.

This work was supported by the Medical Research Council, as part of United Kingdom Research and Innovation (also known as UK Research and Innovation) [MC_UP_1201/30]. For the purpose of open access, the MRC Laboratory of Molecular Biology has applied a CC BY public copyright licence to any Author Accepted Manuscript version arising.

M.A. was funded by the UKRI Medical Research Council (MC_UP_1201/30), A.d.S. was funded by the Wellcome Trust Early Career Award to AdS (227622/Z/23/Z).

## Author contributions

T.H., A.d.S., and M.A. conceived and designed the experiments; T.H. performed all the experiments supervised by A.d.S.; T.H. and A.d.S performed cryo-FIB milling and SEM; P.K. supported data analysis; V.L.H. supported cryo-ET work and data analysis; T.H., A.d.S., and M.A. wrote the manuscript; A.d.S. and M.A. supervised the entire study.

## Declaration of interests

The authors declare no competing interests.

## Supplementary Material

**Supplementary Figure 1.**
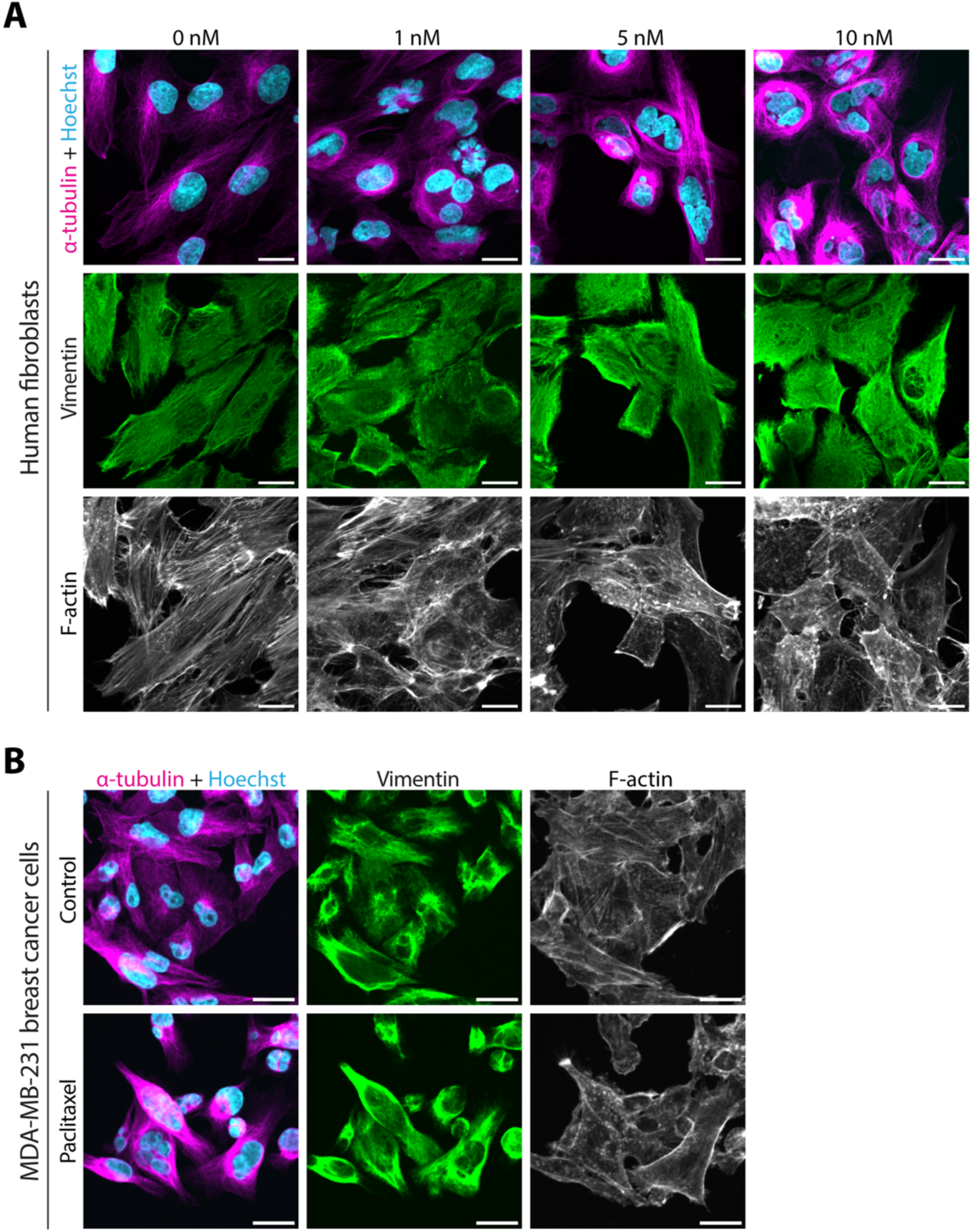
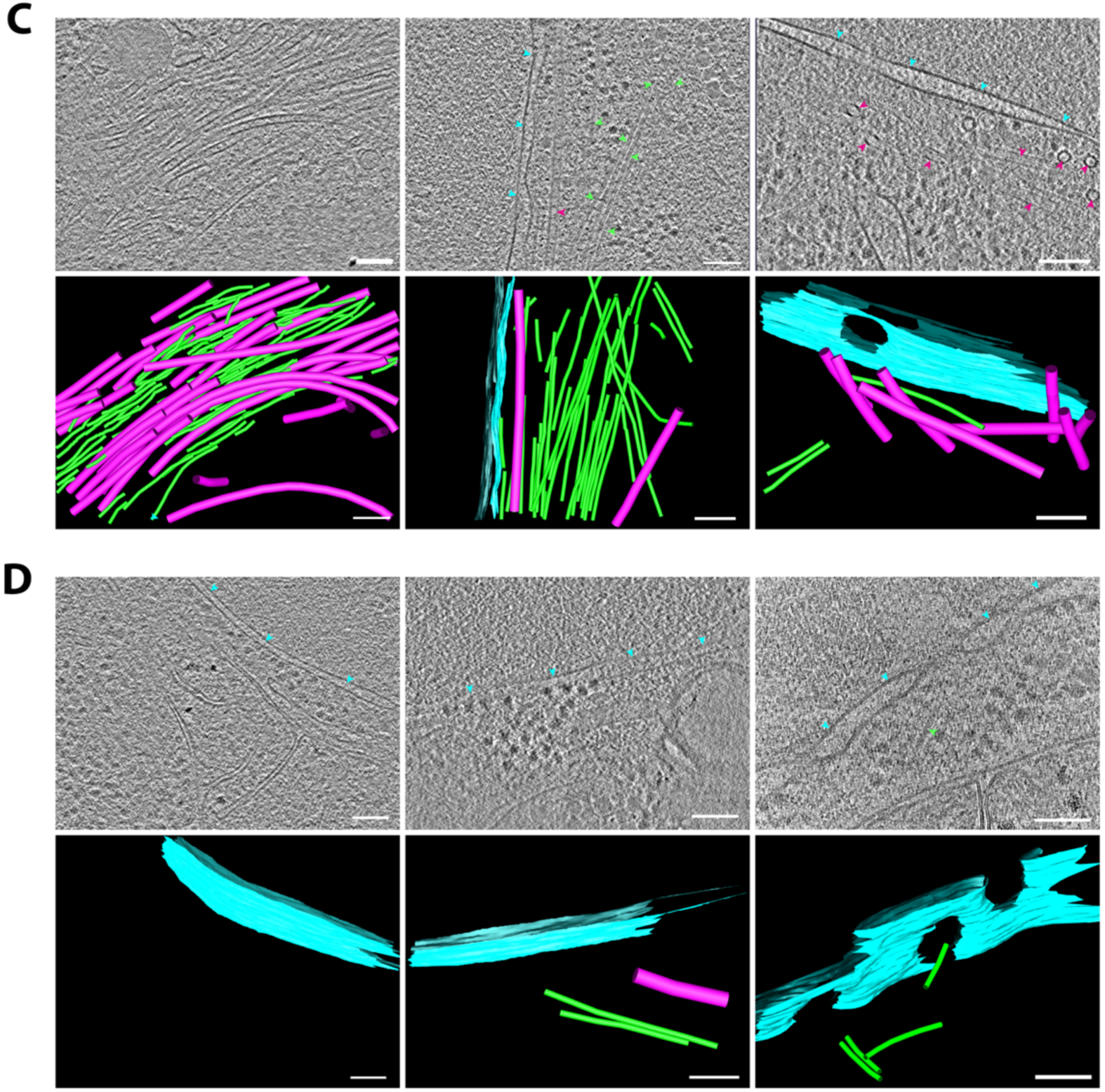
**(A)** Confocal micrographs of human fibroblasts fixed after 16 h incubation in 0, 1, 5, or 10 nM paclitaxel. DNA was stained using Hoechst (cyan), microtubules using α-tubulin immunofluorescence (magenta), vimentin using immunofluorescence (green), and F-actin using phalloidin (white). Scale bars = 20 μm. **(B)** Confocal micrographs of breast cancer MDA-MB-231 cells fixed after 16 h incubation in control media or 5 nM paclitaxel. DNA was stained using Hoechst (cyan), microtubules using α-tubulin immunofluorescence (magenta), vimentin using immunofluorescence (green), and F-actin using phalloidin (white). Scale bars = 20 μm. **(C)** Top panels show 2D slices of reconstructed tomograms from paclitaxel-treated human fibroblasts. Bottom panels show segmentations from these tomograms of microtubules (magenta), vimentin filaments (green), and the NE (cyan), which are marked with arrowheads of the same colour in the tomogram slice. Scale bars = 100 nm. **(D)** As in (C), but with control cells.

**Supplementary Figure 2.**
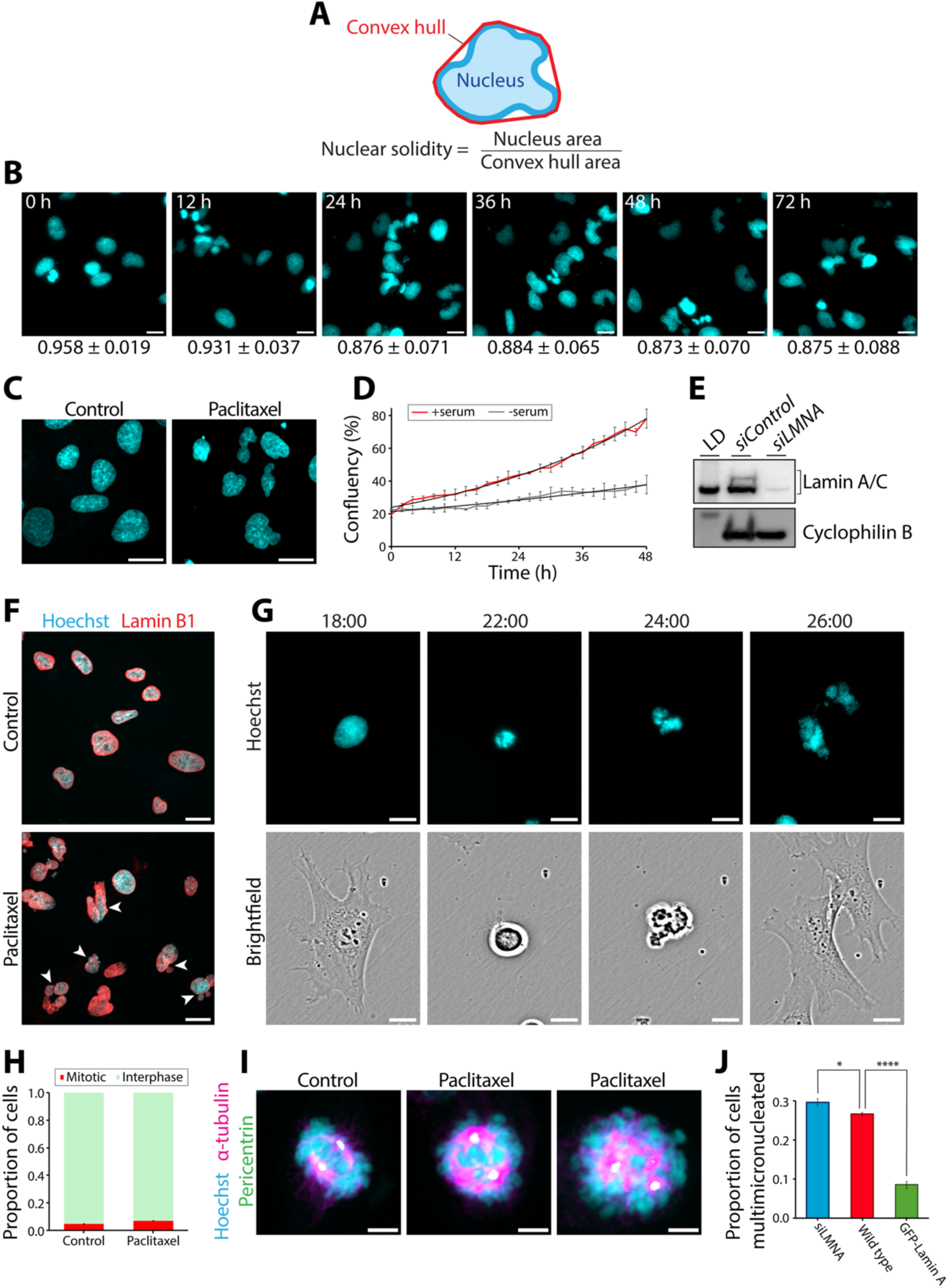
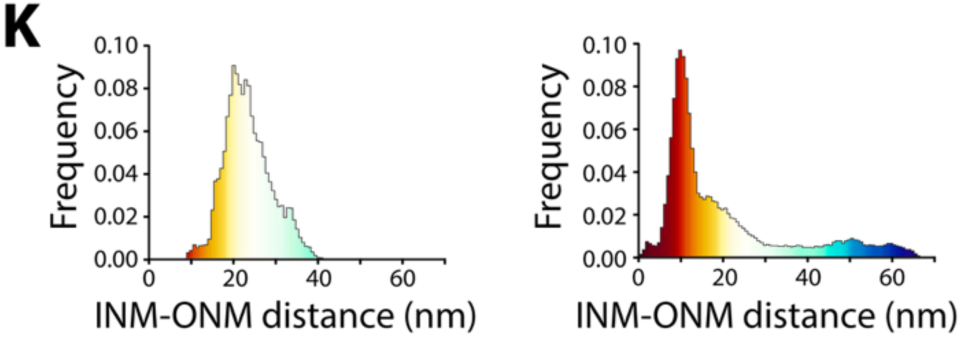
**(A)** Nuclear solidity is used to quantify nuclear deformation and is defined as the nucleus area (blue) divided by the convex hull area of the nucleus^46^ (red). **(B)** Example of images used for nuclear solidity quantification in Figure 2A. Frames from a live-cell movie of Hoechst-stained nuclei 0-72 h after addition of 5 nM paclitaxel are shown, together with the mean nuclear solidity measurements ± the standard deviation from these images. **(C)** Representative images of serum-starved cells following 16 h incubation in control media or 5 nM paclitaxel, that were used for the quantification in Figure 2C. **(D)** Cell confluency of cells cultured over 48 h in complete medium (+serum), or serum-starved medium (-serum). Using non-linear regression to fit exponential growth curves (black) showed that the doubling time was significantly reduced from 28.7 h in complete medium to 58.9 h in serum-starved medium. **(E)** Western Blot confirming knockdown of Lamin A and Lamin C following transfection with siRNA against LMNA (siLMNA), compared to with control siRNA (siControl). Cyclophilin B was used as the loading control. Lane 1 shows protein ladder (LD; 62 kDa for Lamin A/C panel, and 28 kDa for Cyclophilin B panel). **(F)** Confocal micrographs of cells fixed after 24 h incubation in control media or 5 nM paclitaxel, with the nucleus stained using Hoechst (cyan) and Lamin B1 immunofluorescence (red). Multimicronucleated cells are marked with arrowheads. Scale bars = 20 μm. **(G)** Live cell imaging showing a Hoechst-stained cell undergoing mitosis in the presence of 5 nM paclitaxel, 18-26 h after paclitaxel addition. Scale bars = 20 μm. **(H)** Bar graph showing the proportion of cells that were in interphase (green) versus mitosis (red) following 30 h incubation in 5 nM paclitaxel or control media. Mitotic cells were identified by the presence of condensed chromosomes and a mitotic spindle following staining for DNA using Hoechst and microtubules using α-tubulin immunofluorescence. Error bars represent the SEM from three biological repeats (n=3). **(I)** Confocal micrographs of cells fixed after 24 h incubation in control media or 5 nM paclitaxel. Cells were stained for DNA using Hoechst (cyan), microtubules using α-tubulin immunofluorescence (magenta), and centrosomes using pericentrin immunofluorescence (green). Scale bars = 5 μm. **(J)** Bar graph comparing the proportion of wild-type (red), Lamin A/C knockdown (siLMNA – blue), and GFP-Lamin A overexpressing (green) cells that were multimicronucleated following 24 h incubation in 5 nM paclitaxel. Multimicronucleated cells were identified from confocal micrographs of cells stained for DNA using Hoechst. Error bars represent the SEM from three biological repeats (n=3), each with at least 60 cells. Statistical analysis: t-test versus wild type. siLMNA: P=0.0346 (*); GFP-Lamin A: P=4.86 × 10^−5^ (****). **(K)** Histograms showing the ONM-INM distance from the control tomogram (left), and the paclitaxel tomogram (right) in Figure 2G.

**Supplementary Figure 3.**
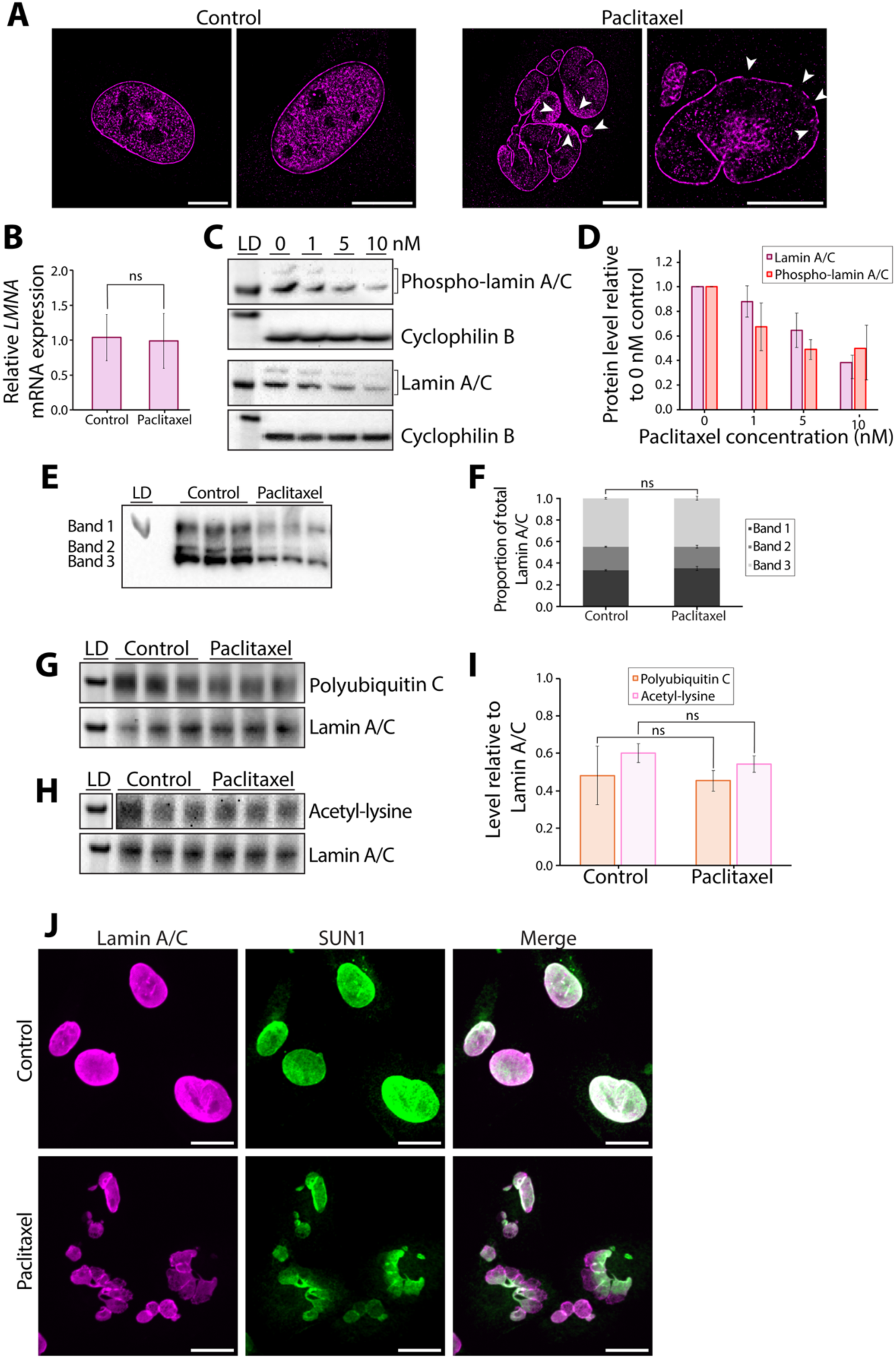
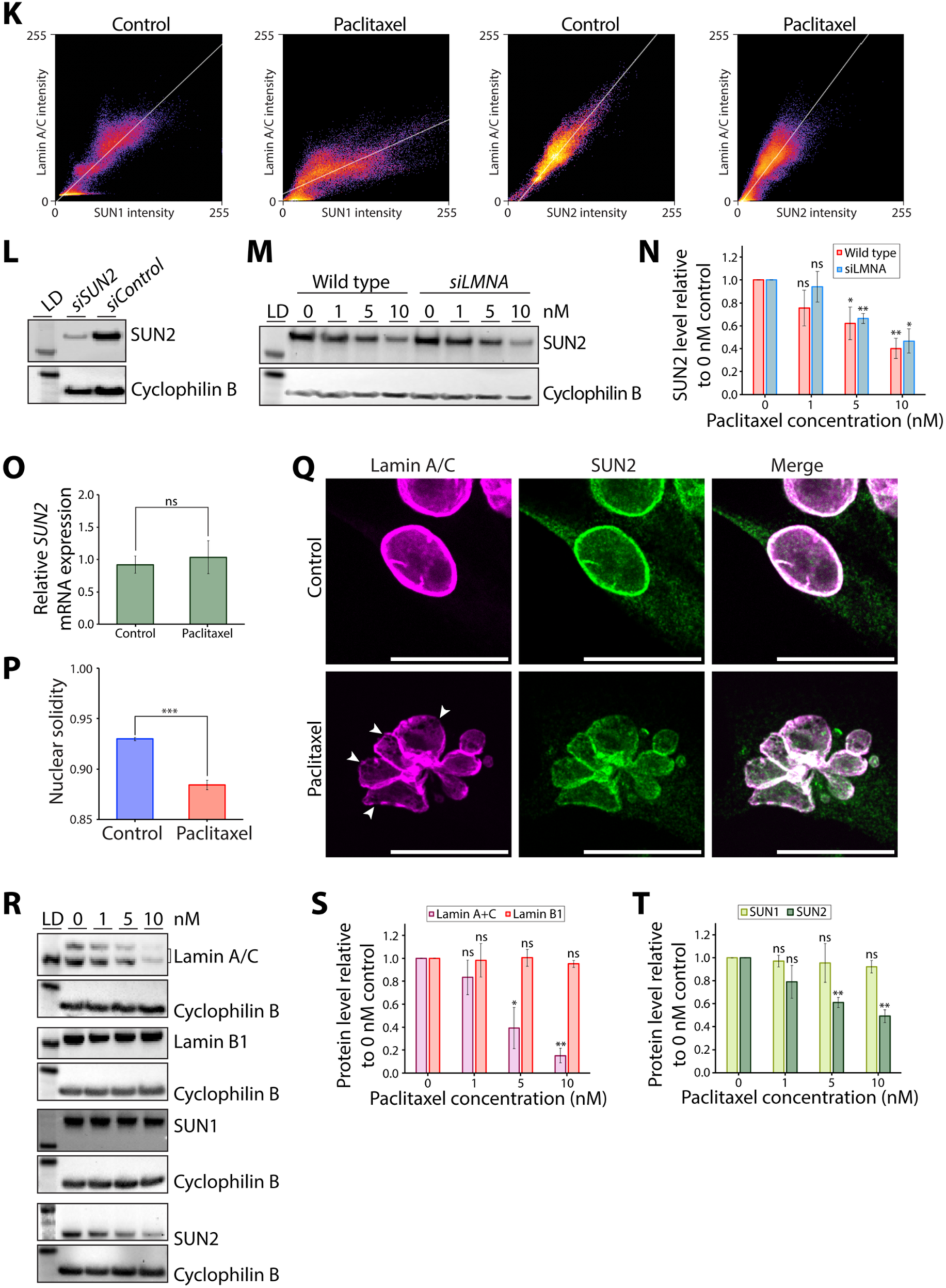
**(A)** STORM images of cells fixed after 16 h incubation in control media or 5 nM paclitaxel. Lamin A/C was stained using immunofluorescence (magenta). Scale bars = 10 μm. Holes in the nuclear lamina are marked with arrowheads. **(B)** Bar chart showing the relative mRNA expression levels of *LMNA* between control cells and cells treated with 5 nM paclitaxel for 24 h, calculated using the 2^−ΔΔCt^ method with GAPDH as the housekeeping gene control. Error bars show the standard deviation from three biological repeats (n=3), each with three technical repeats. Statistical test: t-test, P=0.8742. **(C)** Western blots for Lamin A/C and phospho-Ser404 Lamin A/C of whole cell lysates following 16 h incubation in media containing 0, 1, 5, or 10 nM paclitaxel. Cyclophilin B was used as a loading control. Lane 1 shows protein ladder (LD; 62 kDa for [Phospho]Lamin A/C panels, and 28 kDa for Cyclophilin B panels). **(D)** Quantification of Lamin A/C and phospho-Lamin A/C levels from (C). Each band was normalised to the corresponding Cyclophilin B loading control. Error bars represent the standard deviation from three biological repeats (n=3). **(E)** Western blot for Lamin A/C of whole cell lysates following 16 h incubation in control media or 5 nM paclitaxel. The lysates were run on a Phos-tag gel to separate phosphorylated and non-phosphorylated Lamin A/C. Three biological repeats were used for each condition. Lane 1 shows protein ladder (LD; 62 kDa). **(F)** Quantification of the proportion of total Lamin A/C protein in each of the three bands from (E). Errors bars represent the standard deviation from the three biological repeats (n=3). Statistical test: proportions were first transformed using a centered log-ratio transformation, followed by a MANOVA with Pillai’s trace test, resulting in no significant multivariate effect (P=0.5499). **(G-H)** Top panels: Western blot for **(G)** Polyubiquitin C and **(H)** acetyl-lysine following pull-down of Lamin A/C from whole cell lysates of control cells or cells treated with 5 nM paclitaxel for 16 h. Three biological repeats were used for each condition. Bottom panel: to control for Lamin A/C protein levels, the same membranes were stripped and blotted for Lamin A/C. **(I)** Quantification of Polyubiquitin C from (G) and acetyl-lysine from (H), with each band normalised to Lamin A/C. Error bars represent the standard deviation from three biological repeats (n=3). Statistical analysis: t-test control versus paclitaxel: Polyubiquitin C P=0.7840; acetyl-lysine P=0.1988. **(J**) Confocal micrographs of cells fixed after 16 h incubation in control media or 5 nM paclitaxel. Cells were stained for Lamin A/C (magenta) and SUN1 (green) using immunofluorescence. Scale bars = 20 μm. **(K)** Co-localisation between Lamin A/C and SUN1/SUN2 visualised using scatterplots of Lamin A/C fluorescence intensity versus SUN1/SUN2 fluorescence intensity for each pixel of images in (J) (SUN1) and Figure 3D (SUN2). **(L)** Western Blot confirming knockdown of SUN2 following transfection with siRNA against SUN2 (siSUN2), compared to control siRNA (siControl). Cyclophilin B was used as the loading control. Lane 1 shows protein ladder (LD; 98 kDa for SUN2 panel, and 28 kDa for Cyclophilin B panel). **(M)** Western blots for SUN2 of whole cell lysates from wild-type or Lamin A/C knockdown (siLMNA) cells following 16 h incubation in media containing 0, 1, 5, or 10 nM paclitaxel. Cyclophilin B was used as a loading control. Lane 1 shows protein ladder (LD; 62 kDa for SUN2 panel, and 28 kDa for Cyclophilin B panel). **(N)** Quantification of SUN2 levels from (M). Each band was normalised to the corresponding Cyclophilin B loading control. Error bars represent the standard deviation from three biological repeats (n=3). Statistical analysis: one-sample t-test with null hypothesis mean = 1. Wild type: 1 nM P=0.1112; 5 nM P=0.0445; 10 nM P=0.0070. siLMNA: 1 nM P=0.5221; 5 nM P=0.0053; 10 nM P=0.0132. **(O)** Bar chart showing the relative mRNA expression levels of *SUN2* between control cells and cells treated with 5 nM paclitaxel for 24 h, calculated using the 2^−ΔΔCt^ method with GAPDH as the housekeeping gene control. Error bars show the standard deviation from three biological repeats (n=3), each with three technical repeats. Statistical test: t-test, P=0.5264. **(P)** Bar chart comparing the nuclear solidity of breast cancer MDA-MB-231 cells in control media, or after 16 h incubation in 5nM paclitaxel. Error bars represent the SEM from three biological repeats (n=3), each with more than 100 cells. Statistical test: t-test, P=0.0008. **(Q)** Confocal micrographs of breast cancer MDA-MB-231 cells fixed after 16 h incubation in control media or 5 nM paclitaxel. Cells were stained for Lamin A/C (magenta) and SUN2 (green) using immunofluorescence. Scale bars = 20 μm. **(R)** Western blots for Lamin A/C, Lamin B1, SUN1, and SUN2 of whole cell lysates from breast cancer MDA-MB-231 cells following 16 h incubation in media containing 0, 1, 5, or 10 nM paclitaxel. Cyclophilin B was used as a loading control. Lane 1 shows protein ladder (LD; 62 kDa for Lamin A/C and Lamin B1 panels, 98 kDa for SUN1 and SUN2 panels, and 28 kDa for Cyclophilin B panels). **(S)** Quantification of Lamin A/C and Lamin B1 protein levels from (R). Each band was normalised to the corresponding Cyclophilin B loading control. Error bars represent the standard deviation from three biological repeats (n=3). Statistical analysis: one-sample t-test with null hypothesis mean = 1. Lamin A/C: 1 nM P=0.1981; 5 nM P=0.0279; 10 nM P=0.0019. Lamin B1: 1 nM P=0.8559; 5 nM P=0.9317; 10 nM P=0.1025. **(T)** Quantification of SUN1 and SUN2 protein levels from (R). Each band was normalised to the corresponding Cyclophilin B loading control. Error bars represent the standard deviation from three biological repeats (n=3). Statistical analysis: one-sample t-test with null hypothesis mean = 1. SUN1: 1 nM P=0.4215; 5 nM P=0.6866; 10 nM P=0.1322. SUN2: 1 nM P=0.1263; 5 nM P=0.0043; 10 nM P=0.0041. Ns = non-significant = P>0.05; * P<0.05; ** P<0.01, *** P<0.001.

**Supplementary Figure 4.**
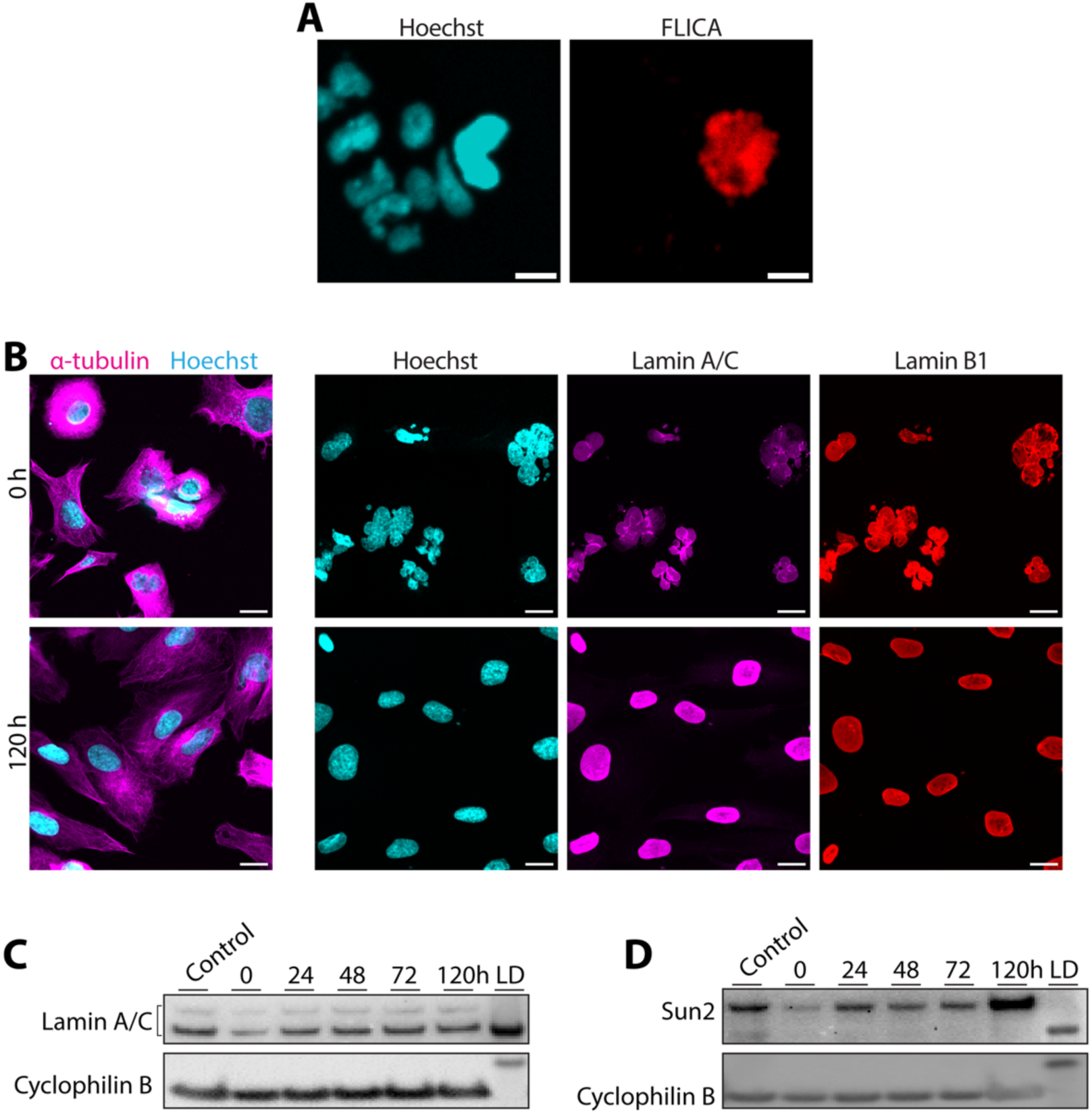
**(A)** Example micrograph used in the cell viability analysis in Figure 5B. Cells were stained for DNA using Hoechst (cyan), and active caspases using FLICA (red). Scale bars = 10 μm. **(B)** Confocal micrographs of cells fixed 0 or 120 h after removal of 5 nM paclitaxel, showing that the microtubule organisation and nuclear lamina recover following paclitaxel removal. Cells were stained for DNA using Hoechst (cyan), and microtubules using α-tubulin immunofluorescence (magenta); or for DNA using Hoechst (cyan), and Lamin A/C (magenta) and Lamin B1 (red) using immunofluorescence. Scale bars = 20 μm. **(C-D)** Western blots for Lamin A/C **(C)** and SUN2 **(D)** of whole cell lysates 0, 24, 48, 72, or 120 h following removal of 5 nM paclitaxel, showing that Lamin A/C and SUN2 protein levels recover following paclitaxel removal. Lane 1 contains untreated cell control. Cyclophilin B was used as a loading control. Lane 7 shows protein ladder (LD; 62 kDa for Lamin A/C and SUN2 panels, and 28 kDa for Cyclophilin B panels).

**Supplementary Table 1.** Details of siRNAs used for knockdowns.

| siRNA | Target mRNA | Source |
| --- | --- | --- |
| siLMNA | *LMNA* | Ambion 4392420 Assay ID s530951 |
| siSUN2 | *SUN2* | Ambion 4392420 Assay ID s24467 |
| siControl | N/A | Ambion 4390844 |

**Supplementary Table 2.** Details of antibodies used for immunofluorescence, Western blotting, and pull-downs.

| **Antibody** | **Dilution (IF – immunofluorescence; WB – Western blot; IP – pull-down)** | **Source** |
| --- | --- | --- |
| Mouse anti-α tubulin | IF: 1:100 | Invitrogen A11126 |
| Rabbit anti-vimentin | IF: 1:200 | Abcam ab92547 |
| Rabbit anti-Lamin B1 | IF: 1:200; WB: 1:2,000 | Abcam ab16048 |
| Mouse anti-Lamin A/C | IF: 1:200; WB: 1:1,000 | Santa Cruz sc7292 |
| Rabbit anti-Lamin A/C | WB: 1:1,000; IP: 1:50 | Abcam ab108595 |
| Mouse anti-phospho-Ser404 Lamin A/C | WB: 1:1,000 | Sigma ABT1387 |
| Rabbit anti-SUN1 | IF: 1:200; WB: 1:2,000 | Sigma HPA008346 |
| Rabbit anti-SUN2 | IF: 1:500; WB: 1:2,000; IP: 1:50 | Abcam ab124916 |
| Rabbit anti-Cyclophilin B | WB: 1:2,000 | Abcam ab16045 |
| Rabbit anti-Polyubiquitin C | WB: 1:500 | Abcam ab104455 |
| Mouse anti-acetyl lysine | WB: 1:500 | Abcam ab22550 |
| Donkey anti-Mouse IgG – Alexa 555 | IF: 1:500 | Abcam ab150110 |
| Donkey anti-Mouse IgG – Alexa 647 | IF: 1:500 | Abcam ab150111 |
| Donkey anti-Rabbit IgG – Alexa 555 | IF: 1:500 | Abcam ab150074 |
| Donkey anti-Rabbit IgG – Alexa 647 | IF: 1:500 | Abcam ab181347 |
| Goat anti-Rabbit IgG - HRP | WB: 1:15,000 | Abcam ab6721 |
| Goat anti-Mouse IgG - HRP | WB: 1:10,000 | Abcam ab97023 |

**Supplementary Table 3.**
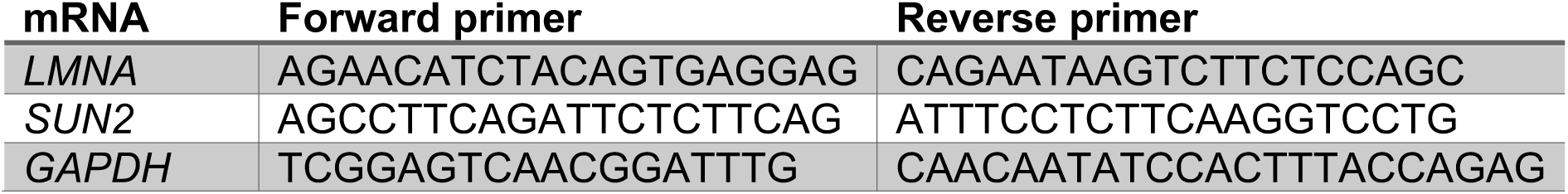
Primer sequences used for RT-qPCR.

## Notes

### Competing Interest Statement

The authors have declared no competing interest.

### Summary of Updates

In this study we investigated the effects of paclitaxel on both healthy and cancerous cells, focusing on alterations in nuclear architecture. Our novel findings show that: 1) Paclitaxel-induced microtubule reorganisation during interphase alters the perinuclear distribution of actin and vimentin. The formation of extensive microtubule bundles, in paclitaxel or following GFP-Tau overexpression, coincides with nuclear shape deformation, loss of regulation of nuclear envelope spacing, and alteration of the nuclear lamina. 2) Paclitaxel treatment reduces Lamin A/C protein levels via a SUN2 dependent mechanism. SUN2, which links the lamina to the cytoskeleton, undergoes ubiquitination and consequent degradation following paclitaxel exposure. 3) Lamin A/C expression, frequently dysregulated in cancer cells, is a key determinant of cellular sensitivity to, and recovery from, paclitaxel treatment. Collectively, our data support a model in which paclitaxel disrupts nuclear architecture through two mechanisms: 1) aberrant nuclear cytoskeletal coupling during interphase, and 2) multimicronucleation following defective mitotic exit. This represents an additional mode of action for paclitaxel beyond its well-established mechanism of mitotic arrest. We thank the reviewers for their time and constructive feedback. We have carefully considered all comments and have carried out a full revision. The updated manuscript now includes additional data showing: 1) Overexpression of microtubule-associated protein Tau causes similar nuclear aberration phenotypes to paclitaxel. This supports our hypothesis that increased microtubule bundling directly leads to nuclear disruption in paclitaxel during interphase. 2) Paclitaxel effects on nuclear shape and Lamin A/C and SUN2 expression levels occur independently of cell division. 3) Reduced levels of Lamin A/C and SUN2 upon paclitaxel treatment occur at the protein level via ubiquitination of SUN2. 4) The effects of paclitaxel on the nucleus are conserved in breast cancer cells. We have also edited our text and added further detail to clarify points raised by the reviewers. We believe that our revised manuscript is overall more complete, solid and compelling thanks to the reviewers comments.

